# Biochar reduces soil thermal conductivity, diffusivity and volumetric heat storage: A global meta-analysis

**DOI:** 10.64898/2026.06.26.734746

**Authors:** Behrouz Gholamahmadi, Damien Beillouin, Kathrin Weber, Lukas Trakal, Ondřej Mašek

## Abstract

Biochar amendments are increasingly applied to improve soil physical functioning and support carbon dioxide removal, but their effects on intrinsic soil thermal properties remain poorly characterised. We conducted the first global systematic meta-analysis of 19 independent studies, 231 control–biochar comparisons, and 529 property-specific effect sizes to test how biochar changes soil heat transfer and storage. Biochar reduced thermal conductivity by 17.6% (95% CI, −22.7 to −12.2), thermal diffusivity by 11.0% (−14.5 to −7.3), and volumetric heat capacity by 8.3% (−12.3 to −4.1). Gravimetric heat capacity showed no significant overall response (+3.3%; −7.6 to 15.4) but was supported by fewer studies. Negative responses were directionally consistent for thermal conductivity, diffusivity, and volumetric heat capacity. Moderator analyses showed that responses were most consistently associated with post-application bulk density and changes in bulk density, while application rate modulated response magnitude and soil texture constrained context dependence. Co-variation among thermal conductivity, thermal diffusivity, and volumetric heat capacity matched expected physical dependencies, indicating coordinated structural reorganisation rather than independent shifts in isolated parameters. These estimates describe intrinsic conductive and storage properties; field-scale soil temperature responses may also be modified by albedo, evaporation, vegetation, and surface energy balance. Improved integration of soil thermal measurements with moisture dynamics, structural changes, and carbon cycling is essential to accurately represent biochar effects in soil and land-surface models.

**Significance Statement:** Biochar is increasingly used to store carbon while improving soils, but its effects on how soils conduct, transmit, and store heat remain poorly understood. By synthesising 19 studies and 529 property-specific effect sizes, we show that biochar consistently reduces soil thermal conductivity, diffusivity, and volumetric heat capacity, primarily through structural changes in bulk density and pore architecture. These findings identify soil thermal behaviour as an overlooked component of biochar–soil interactions, with implications for soil temperature buffering, water–energy coupling, plant and microbial processes, and land-surface and carbon-cycle modelling.

## 1. Introduction

Soil thermal behaviour regulates land-atmosphere energy exchange and underpins biological and biogeochemical processes, including surface temperature dynamics, soil moisture–temperature coupling, microbial activity and plant growth (Hillel, 1998; Bonan, 2019). These processes depend in part on the capacity of soils to conduct, redistribute, and store heat. This capacity is determined by a small set of intrinsic soil thermal properties, including thermal conductivity, thermal diffusivity, and heat capacity.

However, these properties are not fixed material constants: they emerge from the physical organisation of the soil pore system, which governs the continuity of heat transfer pathways and the partitioning of solid, liquid, and gaseous phases (Ochsner et al., 2001; de Vries, 1963; Fatichi et al., 2020). Therefore, soil thermal behaviour is strongly shaped by the soil structure and phase composition. Any process that alters pore geometry, contact networks, or phase distribution can directly modify heat transfer efficiency and bulk thermal storage, rather than acting on intrinsic material conductivity alone. When land management alters soil structure, it can therefore modify soil thermal conductivity, diffusivity, and heat storage capacity.

Despite this structural basis, land-surface and Earth system models commonly use static or weakly responsive parameterisations of soil thermal properties that ignore changes in soil structure induced by management or environmental forcing (Dai et al., 2019; Lawrence et al., 2019; Bonan, 2019). In parallel, experimental studies frequently infer soil thermal behaviour from soil temperature rather than directly quantifying intrinsic thermal properties. Soil temperature integrates multiple external drivers, including atmospheric forcing, soil moisture dynamics, vegetation cover, surface albedo, and surface energy exchange, making it difficult to isolate how land management modifies the intrinsic capacity of soil to conduct, propagate, or store heat. Together, these limitations create a persistent gap between observed soil temperature dynamics and the underlying physical properties that control soil heat transfer.

Biochar represents a substantial structural perturbation of soil physical architecture. This carbon-rich material, produced by pyrolysis of biomass under oxygen-limited conditions, is increasingly used in agricultural management and carbon dioxide removal strategies because of its potential to enhance soil carbon storage and improve soil physical functioning (Woolf et al., 2010; Lehmann & Joseph, 2015; Intergovernmental Panel on Climate Change [IPCC], 2021). Empirical evidence shows that biochar can reduce soil bulk density while increasing porosity, aggregation, water retention, infiltration, and resistance to runoff and erosion (Zhao et al., 2013; Blanco-Canqui, 2017; Pokharel et al., 2020; Schmidt et al., 2021; Gholamahmadi et al., 2023).

These structural changes modify the soil pore network, alter the relative proportions of mineral particles, organic matter, water, and air, and change contact pathways through which heat is transferred. Biochar is therefore expected to affect soil thermal behaviour not only as an added material, but also through the structural reorganisation it induces in the soil matrix. Despite clear evidence that biochar alters soil structure, the effects of biochar on intrinsic soil thermal properties remain inconsistent across studies. Thermal conductivity, thermal diffusivity, volumetric heat capacity, and gravimetric heat capacity are physically interdependent properties.

Yet most studies treat these responses independently, making it difficult to determine whether biochar induces coherent shifts in soil thermal behaviour or context-specific changes in isolated properties. Biochar effects may further vary depending on soil texture, soil moisture, application rate, experimental duration, measurement conditions, and biochar characteristics such as feedstock, pyrolysis temperature, particle size, porosity, ash content, and carbon content. This variability creates uncertainty about how biochar-induced structural reorganisation controls soil heat transport and storage, limiting robust representation of biochar effects in predictive soil and land-surface modelling frameworks.

Here, we address this gap using a global meta-analysis quantifying the effects of biochar on intrinsic soil thermal conductivity, thermal diffusivity, volumetric heat capacity, and gravimetric heat capacity relative to unamended soils, based on 19 independent studies and 231 experimental comparisons. We quantify global response patterns and identify the structural and experimental controls governing variability, including application rate, bulk density change, soil texture, measurement conditions, and biochar material characteristics. Finally, we assess co-variation among thermal properties to determine whether biochar-induced changes are independent shifts in thermal properties or a coordinated reconfiguration of the soil thermal system, providing a mechanistic basis for representing management-induced thermal dynamics in soils.

## 2. Methods

### 2.1 Literature search strategy

A systematic literature search was conducted to identify peer-reviewed experimental studies quantifying the effects of biochar application on intrinsic soil thermal properties. Searches were performed in Web of Science Core Collection and Scopus, covering all years available up to 31 December 2025 and were restricted to articles written in English. Google Scholar was used as a complementary source by screening the first 200 results, sorted by relevance, with screening stopped when no additional eligible records were identified. Reference lists for major review papers and meta-analyses were also screened to maximise retrieval completeness (Blanco-Canqui, 2017; Schmidt et al., 2021).

Search strings combined terms related to biochar, soil, and thermal behaviour. The core query was:

*biochar AND soil AND (“thermal conductivity” OR “thermal diffusivity” OR “heat capacity” OR “volumetric heat capacity” OR “gravimetric heat capacity” OR “thermal resistivity” OR “thermal properties”)*

Additional searches included combinations using related terms such as soil heat transfer, soil thermophysical properties, and thermal behaviour to reduce omissions caused by terminology differences (Supplementary Methods S1). All retrieved records were imported into a reference manager, duplicates were removed, and the remaining records were screened following PRISMA 2020 guidelines. The full screening workflow is shown in Figure S1 and summarised in Table S1.

### 2.2 Study screening and eligibility criteria

Study selection followed a two-stage process consisting of title and abstract screening, followed by full-text eligibility assessment. Eligibility criteria were defined before quantitative extraction to ensure consistency across studies. Studies were retained only when they: (i) experimentally applied intentionally produced biochar to soil under laboratory, column, pot, or field conditions; (ii) included a control treatment without biochar amendment; (iii) directly quantified at least one intrinsic soil thermal property, including thermal conductivity, thermal diffusivity, volumetric heat capacity, gravimetric heat capacity, or thermal resistivity; (iv) reported mean values for both control and biochar-amended treatments; (v) included at least three independent replicates per treatment; and (vi) reported a measure of variability, such as standard deviation, standard error, confidence interval, or sufficient information to reconstruct standard deviations without statistical imputation.

Studies reporting only soil temperature dynamics were excluded because temperature integrates multiple external forcings and does not directly quantify intrinsic thermal transport properties. Studies were also excluded if they used naturally occurring charcoal, historical charcoal, technogenic carbonaceous residues, non-soil substrates, engineered construction media, or hydroponic media to avoid conflating biochar-induced thermal responses with fundamentally different porous materials. No variance imputation was applied for missing dispersion estimates or sample-size information, to avoid artificially inflating precision in a dataset characterised by strong methodological heterogeneity.

### 2.3 Data extraction and dataset construction

For each eligible study, data were extracted across four domains: soil thermal properties, experimental design, soil physical characteristics, and biochar attributes. The extracted response variables included thermal conductivity (*k*), thermal diffusivity (*α*), volumetric heat capacity (*Cv*), gravimetric heat capacity (*Cp*), and thermal resistivity, when reported. For each response variable, means, measures of variability, sample sizes, and units were extracted separately for control and biochar-amended treatments.

The primary experimental unit was defined as the unique experimental contrast within a study, corresponding to a specific combination of soil, biochar treatment, and environmental or measurement conditions sharing a common control. This definition ensures that dependency induced by shared controls and shared experimental settings is explicitly retained in the hierarchical structure of the dataset. When studies reported multiple treatment configurations (e.g. biochar application rates, soil depths, biochar types, moisture conditions, incubation times, or measurement temperatures), each distinct control–biochar contrast was retained as a separate pairwise comparison.

Moderator variables were extracted when reported, prioritising (i) biochar properties, (ii) soil structure, and (iii) environmental conditions. These included biochar application rate, feedstock type, pyrolysis temperature, biochar carbon content, ash content, soil texture, soil organic carbon, soil pH, bulk density, measurement depth, experimental setting, duration after application, moisture condition during measurement, and climate zone. When continuous variables were reported in different units, all values were harmonised before analysis to ensure comparability across studies.

Missing soil texture data (sand, silt, or clay contents) were completed using georeferenced soil databases when coordinates were available. Regional datasets were used where appropriate, and global estimates from SoilGrids250m v2.0 were used when regional data were unavailable, while CSDLv2 was used for georeferenced locations in China (Hengl et al., 2017; Shi et al., 2025). Derived values were used only to complete moderator information and were not treated as primary reported measurements. When numerical values were reported only in figures, data were digitised using plot-extraction procedures and manually checked against the original axes, units, legends, and treatment labels.

Extracted values were cross-checked against the original experimental design, units, and treatment labels before effect-size calculation. This approach follows recent recommendations for transparent extraction of quantitative information from published plots, including vision-assisted and manually verified workflows (Polak & Morgan, 2025).

### 2.4 Effect-size calculation

Effect sizes were calculated for each eligible control–biochar comparison using the natural log response ratio (*lnRR*), which provides a symmetric, dimensionless measure of proportional change suitable for strictly positive thermal properties (Hedges et al., 1999). The *lnRR* was calculated as Eq. 1:

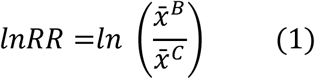

Where 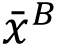 is the mean value of the biochar-amended treatment and 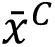 is the mean value of the control treatment. Sampling variances were calculated from treatment and control means, sample sizes, and standard deviations following standard meta-analytic procedures for response ratios (Borenstein et al., 2009; Hedges et al., 1999). When standard deviations were not reported directly, they were reconstructed only from reported standard errors or confidence intervals using standard conversions. Comparisons lacking any extractable measure of variability were excluded; no statistical imputation of missing variance was performed.

Effect sizes were calculated separately for thermal conductivity, thermal diffusivity, volumetric heat capacity, and gravimetric heat capacity. Thermal resistivity values were converted to thermal conductivity using *k = 1/R*, where *R* is thermal resistivity. Converted values were included in the thermal conductivity dataset when corresponding control and biochar treatment means, variability measures, and sample sizes were available. All effect sizes were expressed as percentage change relative to the control for interpretation using Eq. 2:

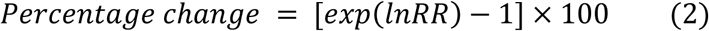

Effect sizes with non-positive sampling variances, arising from reported zero dispersion values, were excluded before model fitting because they would otherwise receive disproportionate statistical weight in inverse-variance meta-analysis. This affected six volumetric heat capacity observations from one study.

### 2.5 Meta-analytic modelling

Separate meta-analyses were conducted for each thermal property using multilevel random-effects models fitted with restricted maximum likelihood. Models included random intercepts for study identity and effect size nested within study, corresponding to a three-level structure with sampling error at the observation level, within-study effect-size heterogeneity, and between-study heterogeneity. This hierarchical structure was used because individual studies frequently reported multiple non-independent observations, allowing the models to account for both within-study non-independence and between-study heterogeneity. The model can be expressed as Eq. 3:

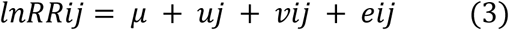

Where *lnRRij* is the log response ratio for comparison *i* in study *j*, *μ* is the overall mean effect, *uj* is the study-level random effect, *vij* is the within-study comparison-level random effect, and *eij* is the sampling error.

Alternative variance structures were compared using AIC, AICc, and BIC: (i) a model with effect-size-level heterogeneity, (ii) a model with study-level heterogeneity only, and (iii) a three-level model with effect sizes nested within studies. The nested three-level structure consistently provided the best fit across all thermal properties and was therefore retained as the primary inferential model (Table S2).

Model outputs were backtransformed from *lnRR* to percentage change relative to the control using Eq. 2. Statistical significance was assessed using 95% confidence intervals and p-values. Heterogeneity was evaluated using Cochran’s Q test and by partitioning variance components between study-level and within-study levels. All analyses were conducted in R version 4.4.0 (R Core Team, 2024). Meta-analytic models were fitted using the metafor package (Viechtbauer, 2010). Data import, processing, visualisation, and spatial mapping were conducted using readxl, dplyr, tidyr, ggplot2, sf, rnaturalearth, and related R packages.

### 2.6 Moderator analysis

Moderator analyses were conducted to identify experimental, soil, and biochar-related factors explaining variation in effect sizes. Candidate moderators were grouped into four mechanistic classes: (i) biochar application characteristics, including application rate and application depth; (ii) biochar material properties, including feedstock, pyrolysis temperature, particle size, carbon content, ash content, pH, and surface area; (iii) soil structural and hydrological properties, including texture, bulk density, porosity, organic carbon, moisture, and water-filled pore space; and (iv) experimental and environmental conditions, including measurement depth, measurement temperature, experimental duration, experiment type, and climate zone.

Each moderator was first evaluated using separate univariate meta-regression models fitted within the same three-level random-effects framework used for the main analysis. Categorical moderators were analysed only when at least two categories had sufficient replication, defined as at least five effect sizes and at least two independent studies per retained category. Variables with many study-specific levels, such as individual experimental design labels, incorporation method, thermal measurement method, and vegetation type, were screened only exploratorily and were not used for primary interpretation because they were closely confounded with study identity. Full moderator-screening results, continuous moderator coefficients, and categorical moderator level estimates are reported in Tables S3–S5.

Moderator effects were assessed using omnibus tests, 95% confidence intervals, and p-values. Results were interpreted according to statistical support, number of effect sizes, number of contributing studies, and mechanistic plausibility. All moderator analyses were univariate; therefore, statistically supported moderators were interpreted as associations rather than definitive causal controls, particularly where moderator variables were incompletely reported or partly confounded with study identity.

### 2.7 Co-variation analysis

Co-variation analysis was used to assess whether biochar-induced changes in soil thermal properties reflect independent shifts in individual parameters or a coordinated reorganisation of the soil thermal system constrained by physical relationships. Because thermal conductivity (k), thermal diffusivity (α), and volumetric heat capacity (Cv) are linked through α = k/Cv, their joint variation provides a thermophysical consistency check on observed responses.

Effect sizes were matched at the level of identical experimental contrasts (study, treatment, and environmental conditions), ensuring comparisons were made within fully equivalent experimental contexts. Co-variation was quantified using correlations among log response ratios (*lnRR*), with Pearson and Spearman coefficients used to assess robustness to non-linearity and outliers. The resulting dataset represents a subset of the full meta-analysis (n = 46–86, depending on property pairing) but preserves strict experimental comparability across all thermal properties.

## 3. Results

### 3.1 Dataset structure and evidence base

The final quantitative synthesis included 19 independent studies and 231 unique pairwise control–biochar comparisons. Thermal conductivity was the most frequently reported property (231 effect sizes from 19 studies), followed by thermal diffusivity (118 effect sizes from 12 studies), volumetric heat capacity (122 effect sizes from 13 studies), and gravimetric heat capacity (58 effect sizes from 6 studies). The evidence base was geographically uneven, with a strong concentration of studies in Asia (n = 14) and fewer studies from Europe (n = 3), North America (n = 1), and Africa (n = 1; Figure S2). Most comparisons were derived from laboratory or controlled experimental settings (n = 195), while pot experiments (n = 8) and field-based measurements (n = 28) remained limited.

This uneven distribution should be considered when interpreting global generalisations, particularly for moderators linked to climate zone, soil texture, and experimental setting. Despite these limitations, the dataset spans a wide range of soil textures, biochar feedstocks, application rates, and thermal measurement conditions, providing sufficient variation to identify first-order structural controls on soil thermal behaviour.

### 3.2 Overall effects of biochar on intrinsic soil thermal properties

Across all studies, biochar induced a coherent and directionally consistent shift in soil thermal behaviour, characterised by reductions in heat transfer and volumetric heat storage. Thermal conductivity showed the strongest response, decreasing by 17.6% relative to unamended controls (95% CI: −22.7 to −12.2%; p < 0.001; 231 effect sizes from 19 studies). Thermal diffusivity decreased by 11.0% (95% CI: −14.5 to −7.3%; p < 0.001; 118 effect sizes from 12 studies), indicating slower propagation of thermal signals through biochar-amended soil (Figure 1). Volumetric heat capacity decreased by 8.3% (95% CI: −12.3 to −4.1%; p < 0.001; 122 effect sizes from 13 studies), indicating a lower capacity to store heat per unit soil volume. In contrast, gravimetric heat capacity showed no significant overall response (+3.3%; 95% CI: −7.6 to 15.4%; p = 0.571; 58 effect sizes from 6 studies), and the evidence base for this property remains too limited to draw firm conclusions.

**Figure 1.**
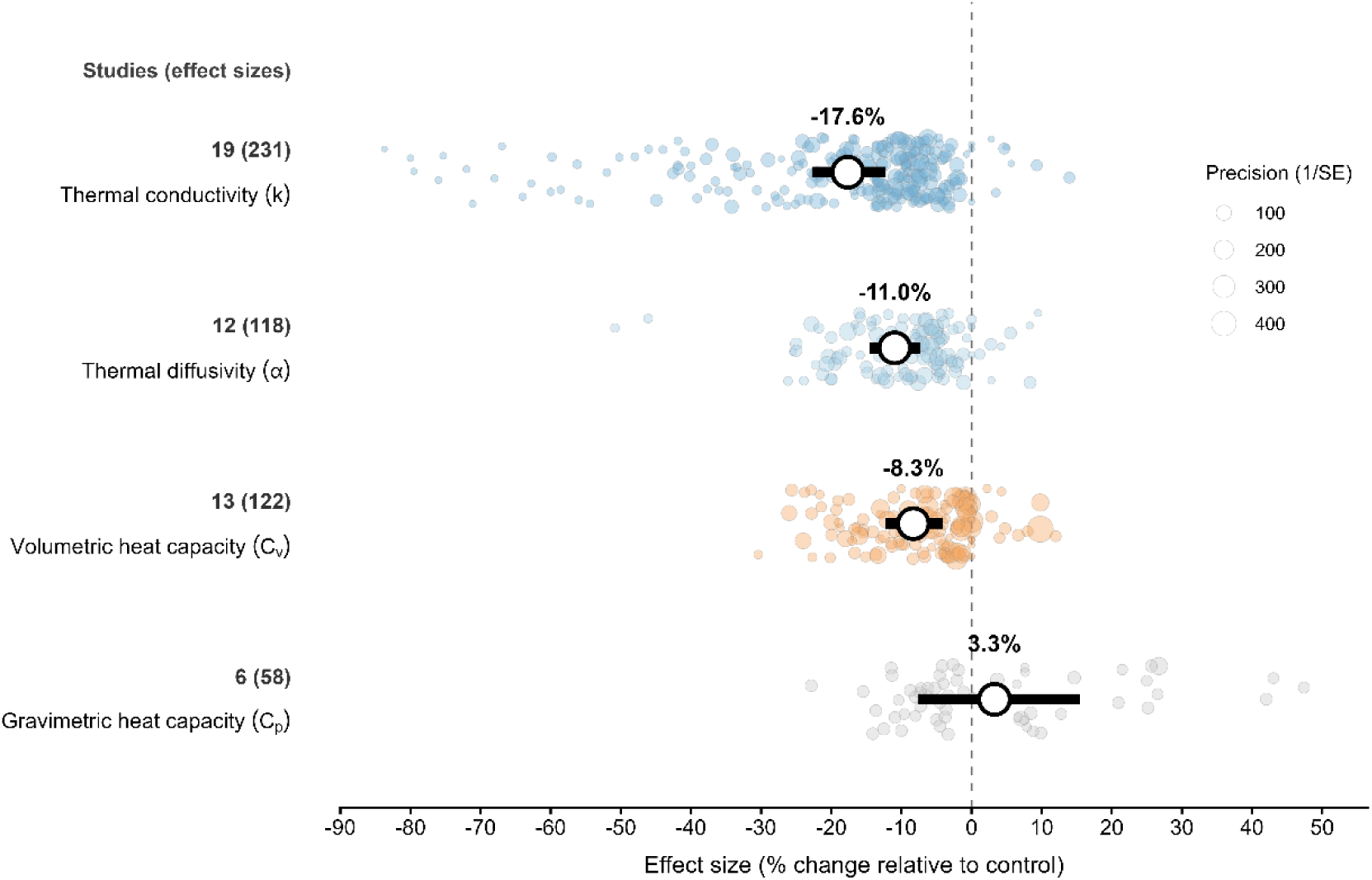
Overall effects of biochar on soil thermal properties. Coloured circles represent individual effect sizes, with circle size scaled by precision (1/SE). Large white circles indicate pooled mean percentage changes estimated using three-level random-effects models, and thick horizontal lines show 95% confidence intervals. Values shown beside each thermal property indicate the number of independent studies and effect sizes, respectively.

The direction of individual effect sizes was also highly consistent for the three transport-or volume-based properties: 95.7% of thermal-conductivity effect sizes, 89.8% of thermal-diffusivity effect sizes, and 85.2% of volumetric-heat-capacity effect sizes were negative. By contrast, only 58.6% of gravimetric-heat-capacity effect sizes were negative, consistent with the weaker and less certain pooled response for this property. The hierarchy of pooled effects therefore followed thermal conductivity > thermal diffusivity > volumetric heat capacity, indicating that biochar most strongly affects heat transfer and heat propagation before volumetric heat storage.

Substantial residual heterogeneity remained for all thermal properties. Total I² was 99.0% for thermal conductivity, 97.0% for thermal diffusivity, 99.2% for volumetric heat capacity, and 98.0% for gravimetric heat capacity. Heterogeneity was partitioned across both study-level and within-study components, supporting the use of the three-level model structure. Cochran’s Q tests were significant for all properties (Q = 6805.8 for k, 2138.6 for α, 9140.9 for Cv, and 2638.2 for Cp; all p < 0.001), indicating substantial unexplained variation among effect sizes. Leave-one-study-out sensitivity analyses confirmed that the direction and statistical significance of the pooled effects were stable for thermal conductivity, thermal diffusivity, and volumetric heat capacity (Table S7). In contrast, gravimetric heat capacity remained non-significant and directionally unstable, consistent with its smaller evidence base.

Small-study-effect diagnostics indicated funnel asymmetry for thermal conductivity (Egger-type test, p < 0.001), whereas tests were not significant for thermal diffusivity (p = 0.367), volumetric heat capacity (p = 0.834), or gravimetric heat capacity (p = 0.719; Table S8). Because funnel asymmetry was significant for thermal conductivity, trim-and-fill sensitivity analyses were performed. No missing effects were imputed (k0 = 0) in either the full effect-size-level analysis or the study-level aggregated analysis, and the adjusted estimates were therefore identical to the unadjusted estimates. The study-level aggregated estimate remained close to the main three-level result (−16.5%), supporting the robustness of the negative thermal-conductivity response while recognising that funnel asymmetry may still reflect heterogeneity, non-independence, or clustered study designs rather than publication bias alone.

### 3.3 Moderator effects

Variation in biochar effects was primarily associated with a small set of structural, application-related, and material controls. Biochar application rate (% w/w) was a significant moderator of thermal conductivity, thermal diffusivity, and volumetric heat capacity. Each one-percentage-point increase in application rate was associated with additional reductions of 1.32% in k (95% CI: −1.48 to −1.16%; p < 0.001), 1.09% in α (95% CI: −1.62 to −0.55%; p < 0.001), and 0.87% in Cv (95% CI: −1.38 to −0.35%; p = 0.001). This dose dependence indicates that soil thermal reorganisation increases with amendment intensity, consistent with a continuous structural perturbation of the soil matrix (Figure 2; Tables S3 and S4).

**Figure 2.**
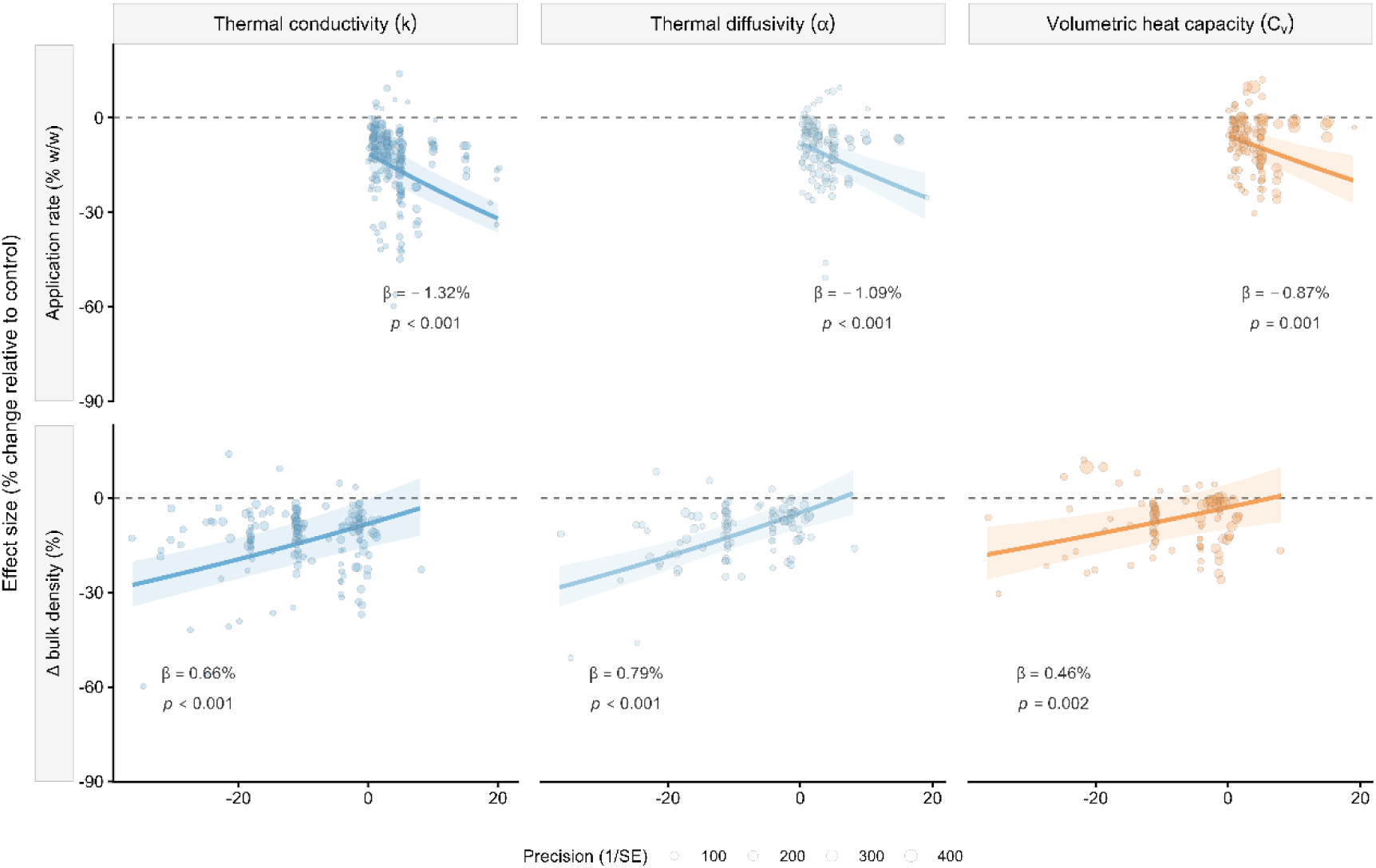
Application-rate and structural controls on biochar effects on soil thermal properties. Panels show relationships between biochar effect sizes and biochar application rate (App. rate; % w/w) or change in bulk density (Δ bulk density; %) for thermal conductivity (k), thermal diffusivity (α), and volumetric heat capacity (Cv). Points show individual effect sizes scaled by precision (1/SE). Lines and shaded bands show fitted meta-regression relationships and 95% confidence intervals from three-level random-effects models. Dashed horizontal lines indicate no effect. Panel β values indicate the change in effect size associated with a one-unit increase in the moderator. For visual clarity, application-rate panels display observations up to 20% w/w, whereas fitted meta-regression slopes were estimated using the full dataset.

Biochar material properties further modulated response magnitude. Feedstock group significantly moderated k (QM = 10.66, p = 0.005) and Cv (QM = 17.52, p < 0.001). Agricultural-residue biochars showed stronger reductions in k (−21.0%; 95% CI: −26.7 to −14.8%) than woody biomass biochars (−7.8%; 95% CI: −17.5 to 3.2%). A similar pattern was observed for Cv, where agricultural-residue biochars reduced Cv by 11.1% (95% CI: −16.3 to −5.5%), while woody biomass biochars showed no clear response. For α, the pyrolysis-temperature group was significant (QM = 7.91, p = 0.019), with a stronger reduction for biochars produced below 450 °C (−13.8%; 95% CI: −17.4 to −9.9%) than for biochars produced above 600 °C (−5.7%; 95% CI: −11.7 to 0.8%). However, the high-temperature group was represented by only seven effect sizes from two studies. Full categorical moderator estimates are reported in Table S5, with omnibus tests summarised in Table S3.

Changes in bulk density represented the clearest structural mediator. Post-application bulk density and change in bulk density were significant moderators for k, α, and Cv. For post-application bulk density, each 1 g cm−3 increase was associated with a 48.0% increase in the response of k (95% CI: 28.4 to 70.7%; p < 0.001), a 27.3% increase in α (95% CI: 9.6 to 47.7%; p = 0.002), and a 35.9% increase in Cv (95% CI: 18.3 to 56.1%; p < 0.001). Change in bulk density also showed significant positive slopes for k, α, and Cv, with each one-percentage-point increase in bulk-density change associated with increases of 0.66% for k (95% CI: 0.39 to 0.92%; p < 0.001), 0.79% for α (95% CI: 0.49 to 1.08%; p < 0.001), and 0.46% for Cv (95% CI: 0.16 to 0.76%; p = 0.002). Because biochar generally reduced bulk density, these positive slopes indicate that stronger reductions in bulk density were associated with stronger reductions in heat transfer, heat propagation, and volumetric heat storage.

Soil texture and environmental context further constrained the response. Soil texture significantly moderated Cv (QM = 23.56, p < 0.001), with the strongest reductions observed in clay loam soils (−17.0%; 95% CI: −21.7 to −12.0%) and sandy loam soils (−9.6%; 95% CI: −13.9 to −5.1%), while silt loam soils showed no clear response (−0.2%; 95% CI: −4.9 to 4.7%). Climate zone moderated k (QM = 4.37, p = 0.037) and Cv (QM = 6.33, p = 0.012), with stronger reductions in subtropical than temperate conditions. Soil moisture significantly moderated Cv (0.18% change per one-percentage-point increase in soil moisture; 95% CI: 0.08 to 0.28%; p < 0.001), but not k or α. Measurement temperature was a significant moderator for k and α, with higher measurement temperature associated with stronger reductions in both properties. Experimental duration showed no clear moderator effect, and application depth was too inconsistently reported to support robust quantitative interpretation.

Overall, these moderators support a structural interpretation: biochar effects on soil thermal behaviour are primarily associated with structural modification of the soil matrix, with application rate setting the perturbation magnitude and soil texture, moisture state, measurement conditions, and biochar properties constraining the response envelope. Because all moderator analyses were univariate, these associations should be interpreted cautiously; correlated moderators such as feedstock, pyrolysis temperature, application rate, geography, and study identity may partly confound individual moderator effects (Tables S3–S5).

### 3.4 Co-variation among thermal properties

The joint responses of thermal conductivity and volumetric heat capacity provide a quantitative consistency check on the thermophysical coupling governing soil heat transport. The observed reduction in thermal diffusivity (−11.0%) closely matched the value predicted from the coupled responses of thermal conductivity and volumetric heat capacity (−10.1%, computed on the *lnRR* scale as *exp[lnRR(k) − lnRR(Cv)] − 1*), yielding a residual deviation of approximately 0.9 percentage points. This agreement indicates that biochar-induced changes in thermal diffusivity are largely emergent from coordinated shifts in heat transfer and heat storage properties, rather than representing an independent response (Figure 3; Table S6).

**Figure 3.**
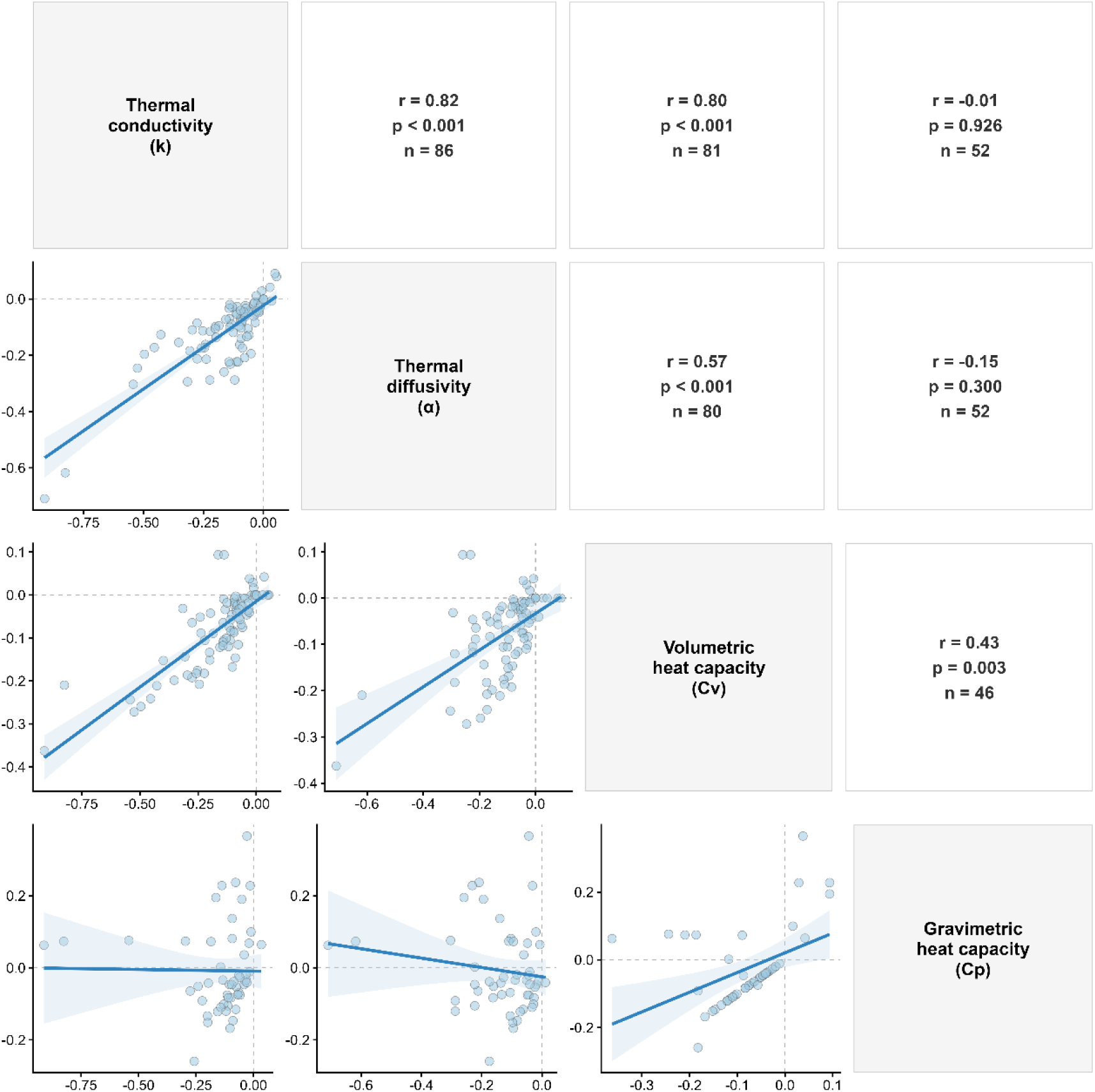
Co-variation among biochar-induced changes in intrinsic soil thermal properties. Lower panels show pairwise relationships among log response ratios for thermal conductivity (k), thermal diffusivity (α), volumetric heat capacity (Cv), and gravimetric heat capacity (Cp). Points represent comparison-level effect sizes where both properties were reported for the same experimental contrast. Blue lines and shaded bands indicate linear relationships and 95% confidence intervals. Upper panels report Pearson correlation coefficients, p-values, and pairwise sample sizes. Dashed lines indicate zero effect.

Biochar-induced changes in soil thermal properties showed a strongly coupled response structure for heat-transfer and volumetric heat-storage variables, contrasting with a weaker and less certain response for mass-normalised heat capacity. Thermal conductivity showed strong positive correlations with thermal diffusivity (Pearson r = 0.82, p < 0.001; Spearman ρ = 0.74, p < 0.001; n = 86) and volumetric heat capacity (Pearson r = 0.80, p < 0.001; Spearman ρ = 0.78, p < 0.001; n = 81). Thermal diffusivity and volumetric heat capacity were also positively correlated, although less strongly (Pearson r = 0.57, p < 0.001; Spearman ρ = 0.58, p < 0.001; n = 80).

In contrast, gravimetric heat capacity was not significantly correlated with thermal conductivity (Pearson r = −0.01, p = 0.926; Spearman ρ = 0.26, p = 0.058; n = 52) or thermal diffusivity (Pearson r = −0.15, p = 0.300; Spearman ρ = −0.08, p = 0.555; n = 52), although it was moderately associated with volumetric heat capacity (Pearson r = 0.43, p = 0.003; Spearman ρ = 0.53, p < 0.001; n = 46). This divergence indicates that biochar primarily modifies soil thermal behaviour through structural and volumetric reorganisation of the soil matrix, while evidence for consistent changes in intrinsic mass-normalised heat capacity remains limited.

These results confirm that biochar effects on soil thermal properties are not independent. Instead, they reflect a coordinated shift involving heat transfer, heat propagation, and volumetric heat storage. The weaker coupling of Cp with k and α supports the interpretation that biochar primarily modifies soil thermal behaviour through changes in soil structure and volumetric composition rather than through consistent changes in intrinsic mass-based heat capacity.

## 4. Discussion

### 4.1 Biochar effects on soil thermal properties are structurally driven

Soil thermal theory predicts that heat transfer and storage are governed by the relative proportions and spatial arrangement of mineral particles, organic matter, water, and air within the pore system, with solid–solid contact networks and phase continuity providing the primary pathways for heat conduction (de Vries, 1963; Campbell et al., 1994; Ochsner et al., 2001; Lu et al., 2014; Tong et al., 2016). Under this framework, any amendment that reduces bulk density, modifies pore architecture, or alters volumetric phase composition is expected to reduce thermal conductivity and, through it, thermal diffusivity and volumetric heat capacity.

Biochar satisfies these conditions: its incorporation reduces bulk density, increases porosity, and introduces a carbonaceous phase with intrinsic thermal conductivity substantially lower than most mineral soil constituents (Patwa et al., 2022). Individual experiments have reported reductions in k and α consistent with these mechanisms (Zhang et al., 2013; Zhao et al., 2016; Usowicz et al., 2016, 2020), but studies remained small, geographically concentrated, and methodologically heterogeneous. Whether the direction and hierarchy of biochar effects were globally consistent, and whether changes in individual thermal properties were coupled or independent, remained open empirical questions.

The present synthesis resolves these questions quantitatively. The pooled reduction in thermal conductivity (−17.6%) exceeded that of thermal diffusivity (−11.0%), which in turn exceeded that of volumetric heat capacity (−8.3%), a hierarchy consistent with the physical relationship α = k/Cv and with the expectation that amendments disrupting solid–solid contact would most strongly affect conductive pathways. The co-variation analysis reinforces this interpretation: biochar-induced changes in k, α, and Cv were strongly and positively correlated, indicating that these properties responded as a physically coupled system rather than as independent outcomes.

The internal thermophysical consistency of the observed responses provides additional support: the predicted change in thermal diffusivity derived from the coupled responses of k and Cv (−10.1%) closely matched the observed pooled estimate (−11.0%), with a residual deviation of less than one percentage point. This agreement would not be expected if the three properties were responding to independent or arbitrary drivers and confirms that biochar modifies soil heat transport through coordinated structural reorganisation rather than through isolated changes in individual thermal parameters.

The contrasting response of gravimetric heat capacity provides an important constraint on this interpretation. Unlike volumetric heat capacity, which is expressed per unit volume and is therefore sensitive to bulk density, porosity, and water content, gravimetric heat capacity is expressed per unit mass. Biochar commonly has a much lower bulk density than mineral soil, meaning that a given mass of biochar can occupy a disproportionately large volume within the soil matrix. This can substantially alter volumetric composition and pore architecture without producing a consistent change in mass-normalised heat capacity. The weak and statistically non-significant Cp response should therefore be interpreted cautiously: it likely reflects both the mass-based definition of Cp and the smaller evidence base for this property, rather than evidence that biochar has no influence on soil heat storage.

### 4.2 Context dependence: application rate, soil texture, moisture and biochar properties

Although the overall direction of biochar effects was consistent for k, α, and Cv, the magnitude of response varied substantially among studies. The clearest controls were biochar application rate and soil structural change. Higher application rates were associated with stronger reductions in thermal conductivity, thermal diffusivity, and volumetric heat capacity, indicating that biochar effects scale with the degree of physical alteration imposed on the soil. Importantly, these rate effects should be interpreted as evidence of dose-dependent structural change rather than as a universal recommendation for higher biochar application rates. Rate-dependent changes in soil thermal behaviour have also been reported in individual biochar experiments, including studies showing that increasing biochar addition reduced thermal conductivity and modified soil temperature dynamics (Zhang et al., 2013; Khaledi et al., 2023).

This dose dependence is consistent with a mechanism in which increasing biochar inputs progressively modify the volumetric composition of the soil matrix, reduce mineral contact continuity, and alter the balance among solid, water-filled, and air-filled phases. Bulk-density variables provided the strongest structural evidence in the moderator analysis. Because biochar generally reduces bulk density, the positive slopes for post-application bulk density and bulk-density change indicate that stronger reductions in bulk density were associated with stronger reductions in heat transfer, heat propagation, and volumetric heat storage. This does not imply that bulk density alone controls the response, but it identifies bulk-density change as a useful integrative indicator of biochar-induced structural reorganisation.

This supports a structural mechanism in which biochar increases pore volume and matrix heterogeneity, disrupts mineral contact continuity, increases the tortuosity of conductive pathways, and enhances the relative contribution of air-filled pores. However, the thermal consequence of added pore space depends strongly on whether pores are air-filled or water-filled. In the present moderator analysis, this moisture signal was detectable for Cv but not for k or α, suggesting that reported moisture conditions captured variation in volumetric heat storage more clearly than variation in conductive or diffusive heat transfer. The magnitude of this response is likely modulated by soil moisture conditions, because water and air differ strongly in their capacity to conduct and store heat, and soil thermal conductivity models consistently identify water content, porosity, bulk density and texture as dominant controls (Lu et al., 2014; Tong et al., 2016; Xie et al., 2020).

Soil texture further constrained the response, particularly for Cv. Stronger reductions were observed in clay loam and sandy loam soils, whereas silt loam soils showed no clear response. This weaker response in silt loam may reflect its intermediate pore-size distribution and lower sensitivity to additional disruption of conductive pathways under the measured conditions. In sandy soils, biochar may more strongly disrupt grain-to-grain contact and quartz-dominated conductive pathways, whereas in finer-textured soils, changes in water retention and water-mediated thermal bridges may be more important. Thus, the same biochar addition may reduce heat transfer through different physical pathways depending on soil texture and moisture state.

This interpretation is consistent with evidence that biochar effects on water retention are texture-dependent and can be stronger in coarse-textured soils when biochar pore volume and particle size increase soil water storage capacity, with biochar particle size and surface hydrophobicity acting as direct controls on soil hydraulic behaviour (Wang et al., 2019; Razzaghi et al., 2020; Edeh & Mašek, 2022; Gholamahmadi et al., 2025a, 2025b). However, the texture-specific interpretations should be treated cautiously because texture effects were partly confounded with study identity, moisture conditions, and limited replication within some texture classes.

Biochar material properties also contributed to context dependence. Agricultural-residue biochars were associated with stronger reductions in k and Cv than woody biomass biochars, and lower-temperature biochars were associated with stronger reductions in α than high-temperature biochars (Zhao et al., 2021). This pattern is plausible because feedstock selection and pyrolysis conditions strongly regulate biochar physicochemical properties, including carbon and ash content, surface chemistry, density, surface area, pore architecture, and water interactions (Weber & Quicker, 2018; Gholamahmadi, 2026; Zhao et al., 2013). Direct 4D synchrotron microtomography shows that biochar porosity, pore volume, and pore connectivity vary substantially with both feedstock and pyrolysis temperature (Edeh et al., 2023), while molecular-scale biochar models with controlled porosity provide a route to linking production conditions to pore-network geometry (Ngambia et al., 2024).

Biochar ageing and physical degradation may further modify these interactions because soil exposure, wetting–drying, abrasion, fragmentation, and pore alteration can change biochar wettability, porosity, surface area, and water-holding behaviour over time (Ivanova et al., 2023; Barbosa et al., 2025). These material effects could modify soil thermal responses by altering the balance among low-conductivity carbonaceous particles, air-filled pore space, water-filled pore space, and mineral contact pathways. However, they remain indicative rather than definitive because particle density, pore-size distribution, ash composition, moisture state, ageing status, and intrinsic biochar thermal conductivity were rarely reported and often covaried with study design.

The intrinsic thermal conductivity of biochar itself is likely an important but underreported mechanism. Patwa et al. (2022), for example, reported thermal conductivity values of approximately 0.10–0.13 W m^−1^ K^−1^ for produced biochars across pyrolysis temperatures, substantially lower than many mineral soil constituents. This supports the interpretation that porous carbonaceous material can reduce conductive heat pathways when incorporated into soil. However, because most studies did not report intrinsic biochar thermal conductivity, this property could not be included as a quantitative moderator. Future studies should therefore report not only feedstock and pyrolysis temperature, but also biochar bulk density, particle density, porosity, particle-size distribution, moisture state, and intrinsic thermal conductivity, because these properties are needed to separate material effects from soil-structural effects.

### 4.3 Practical implications for soil management and modelling

The results have practical implications for both biochar management and soil thermal modelling. From a management perspective, application rate and bulk-density change emerge as key variables linking biochar use to soil thermal response. Higher biochar application rates produced stronger reductions in k, α, and Cv, but these effects should not be interpreted as universally beneficial or detrimental. Lower thermal conductivity and diffusivity may improve thermal buffering by slowing heat propagation through the soil profile, but the agronomic consequence will depend on climate, season, crop sensitivity, soil moisture, and placement depth.

In cooler or early-season conditions, reduced heat propagation may delay soil warming and potentially affect germination or seedling emergence, whereas in hot or drought-prone environments, it may help dampen thermal extremes in the rhizosphere. Therefore, biochar application should be evaluated not only in terms of carbon storage or water retention, but also in terms of how it modifies the soil thermal regime under local pedoclimatic conditions. This requires considering application rate together with biochar type, incorporation depth, soil texture, moisture regime, and crop or ecosystem objective.

For experimental design and reporting, the findings indicate that post-application bulk density, application rate, soil moisture, and measurement depth should be treated as core variables rather than auxiliary information. Without these measurements, it is difficult to determine whether observed thermal responses arise from biochar material properties, soil structural reorganisation, moisture redistribution, or measurement conditions. This is particularly important because the same biochar dose can produce different thermal outcomes depending on texture, moisture status, and the extent of incorporation into the soil matrix.

For modelling, the results provide a concrete entry point for representing biochar effects in soil thermal frameworks. Land-surface models typically parameterise k from bulk density, porosity, texture, and water content (Lu et al., 2014; Tong et al., 2016; Dai et al., 2019; Lawrence et al., 2019). Since post-application bulk density was the strongest moderator of biochar-induced changes in k, α, and Cv, the most tractable approach is to update these physical inputs, rather than add biochar-specific thermal parameters, to reflect post-application structural conditions. Representing biochar effects through measurable changes in bulk density, porosity, and moisture retention, rather than as a fixed thermal offset, would allow existing pedotransfer functions to propagate biochar-induced structural changes into soil heat transfer and storage predictions without requiring new model architectures.

### 4.4 Relevance for biochar carbon removal and field-scale energy balance

Biochar is widely discussed as a durable carbon dioxide removal pathway because a fraction of its carbon can persist in soils over long timescales, depending on feedstock, pyrolysis conditions, environmental context, and post-application fate (Woolf et al., 2010; Crombie et al., 2013; Lehmann & Joseph, 2015; IPCC, 2021; Weng & Cowie, 2025). Most assessments of biochar carbon persistence focus on chemical recalcitrance, aromatic carbon structure, mineral interactions, and physical protection from decomposition. The present synthesis suggests that biochar may also influence the soil physical environment through a less explored thermal pathway: the modification of heat transfer, heat propagation, and volumetric heat storage.

Our estimates describe intrinsic, conduction-and storage-related properties measured largely under controlled moisture and density conditions. They do not capture field-scale processes such as surface albedo and shortwave absorption, evaporative and latent-heat effects, advective or vapour-phase heat transport, or freeze–thaw dynamics. Because these processes can act in opposing directions, lower intrinsic conductivity may amplify, offset, or even reverse field-scale thermal outcomes depending on surface cover, incorporation depth, moisture status, vegetation, and seasonal temperature gradients. The observed reductions in thermal conductivity and diffusivity should therefore be interpreted as changes in the conductive and storage capacity of the soil matrix, not as direct evidence for additional carbon removal.

These thermal effects should be interpreted as implications rather than direct evidence for enhanced carbon persistence. Thermal buffering may help stabilise rhizosphere temperature, reduce thermal stress on microbial activity, influence temperature-sensitive greenhouse-gas dynamics, and affect seedling emergence through altered soil temperature profiles (Li et al., 2020; Wang et al., 2023; Xu et al., 2023). However, soil moisture, substrate availability, oxygen supply, microbial community composition, vegetation feedbacks, soil texture, and surface energy balance also regulate carbon and greenhouse-gas responses. Therefore, lower thermal conductivity should not be interpreted as a quantified carbon-persistence or greenhouse-gas mitigation effect. Rather, it identifies a physical process that should be considered alongside chemical and hydrological mechanisms when evaluating biochar effects on soil carbon outcomes.

This distinction is particularly important at the field scale, where intrinsic soil thermal properties interact with surface radiative processes. Biochar can darken the soil surface and reduce albedo or reflectance when it remains exposed or concentrated near the surface, potentially increasing absorbed radiation (Genesio et al., 2012; Meyer et al., 2012; Verheijen et al., 2013; Zhang et al., 2013; Usowicz et al., 2016). Under such conditions, heat flux may not decline in proportion to thermal conductivity because field-scale heat transfer depends on both soil thermal conductivity and the imposed temperature gradient. Thus, lower intrinsic thermal conductivity in biochar-amended soil does not necessarily imply lower surface heating or lower vertical heat flux under all field conditions. The net outcome will depend on incorporation depth, vegetation cover, residue cover, soil moisture, albedo, and seasonal temperature gradients. Field-scale studies should therefore evaluate intrinsic thermal properties together with surface energy balance, rather than assuming that lower conductivity directly translates into lower soil heating.

Previous syntheses show that biochar can improve water retention, infiltration, aggregation, and erosion resistance (Blanco-Canqui, 2017; Razzaghi et al., 2020; Gholamahmadi et al., 2023; Ghorbani & Amirahmadi, 2024), and these hydrological and structural effects can influence the physical protection and redistribution of soil organic carbon. The present synthesis adds a complementary thermal dimension by showing that biochar also modifies the physical environment in which soil carbon processes occur. However, the CDR relevance of this thermal pathway remains an emerging hypothesis rather than a quantified removal-crediting mechanism. Current evidence is insufficient to assign a direct carbon-persistence benefit to the observed thermal changes (Gholamahmadi & Kammann, 2026).

A further untested implication concerns biochar permanence itself. Current permanence models commonly include temperature-dependent mineralisation rates, so soil temperature can influence the estimated fraction of biochar carbon remaining over a century (Woolf et al., 2021; Azzi et al., 2024). By reducing thermal conductivity and diffusivity, biochar could theoretically moderate part of the temperature regime that influences its own persistence. This possibility remains qualitative and unresolved: thermal buffering affects temperature amplitude, whereas permanence models are more sensitive to mean soil temperature and model-structure assumptions. Testing this feedback would require coupling intrinsic thermal-property measurements with soil temperature profiles and permanence modelling across contrasting climates.

### 4.5 Limitations and future research

Several limitations should be considered when interpreting these findings. First, the evidence base remains relatively small and unevenly distributed across thermal properties. Thermal conductivity was represented by the largest number of studies and effect sizes, whereas gravimetric heat capacity was reported in only six studies. This imbalance is consistent with the broader experimental literature, where thermal conductivity is more commonly measured than heat capacity or diffusivity in biochar-amended soils (Usowicz et al., 2016; Zhao et al., 2016; Khaledi et al., 2023). This imbalance limits inference for Cp and partly explains its directional instability in sensitivity analyses.

Second, the geographic distribution of studies was uneven, with a strong concentration in Asia and fewer studies from Europe, North America, and other regions. Most observations were also derived from laboratory or controlled experiments, while field-scale measurements remained limited. This restricts direct extrapolation to agricultural and restoration systems where vegetation cover, seasonal moisture dynamics, solar radiation, surface residues, and management history influence soil thermal regimes. The results should therefore be interpreted as a synthesis of the available experimental evidence rather than as a globally resolved field-scale prediction.

Third, differences in experimental and measurement conditions introduced substantial heterogeneity. Studies varied in soil moisture status, measurement temperature, measurement depth, experimental duration, application rate, incorporation method, and thermal measurement technique. These differences are not only methodological sources of variation; they are part of the physical system controlling soil heat transfer. This is consistent with soil thermal models and experiments showing strong sensitivity of thermal conductivity to water content, bulk density, porosity and texture (Lu et al., 2014; Tong et al., 2016; Xie et al., 2020).

Moisture data were sufficient to test as a moderator for some properties, but reporting was too inconsistent to isolate moisture-state effects across the full dataset. Future studies should therefore report thermal conductivity, thermal diffusivity, volumetric and gravimetric heat capacity, bulk density, porosity, water content, measurement temperature, measurement depth, application rate, incorporation depth, residence time, and soil texture in a standardised way. Both volumetric and gravimetric heat capacity should be reported together with bulk density, water content, and biochar particle density, so that mass-based and volume-based thermal responses can be distinguished mechanistically.

Fourth, all moderator analyses were conducted univariately. This is standard practice in meta-analysis with limited data, but it means that moderators may be correlated with each other and with study identity. For example, apparent climate-zone effects may partly reflect geographic differences in study design or application rate, and feedstock effects may covary with pyrolysis temperature or biochar particle structure. Moderator results should therefore be interpreted as plausible associations that identify candidate mechanisms, not as formal variance partitioning or definitive causal attribution.

Fifth, biochar material properties were incompletely reported. Feedstock, pyrolysis temperature, particle size, surface area, carbon content, and ash content were available for some studies, but intrinsic biochar thermal conductivity, particle density, porosity, moisture state, and particle-size distribution were rarely reported. This gap is important because available measurements suggest that biochar can have substantially lower thermal conductivity than mineral soil constituents, but values depend on feedstock, pyrolysis conditions, porosity and moisture state (Patwa et al., 2022). These variables are important because the final soil thermal response depends on both the thermal behaviour of the biochar phase and the structural reorganisation of the surrounding soil matrix. Longer-term studies are also needed to determine whether biochar effects on soil thermal properties persist, weaken, or change direction as biochar surfaces age, hydrophobicity changes, pores become colonised or clogged, and soil structure reorganises after application. This is particularly important because freshly produced biochar may behave differently from aged biochar that has undergone wetting–drying cycles, microbial colonisation, mineral coating, or partial pore blockage in soil.

Finally, this meta-analysis focused on intrinsic soil thermal properties rather than full field-scale energy balance. Albedo, surface temperature, sensible and latent heat fluxes, vegetation cover, soil respiration, and greenhouse-gas fluxes were generally not measured together with thermal conductivity or heat capacity. Future field studies should therefore couple biochar thermal-property measurements with soil moisture, temperature profiles, surface albedo, evapotranspiration, soil respiration, microbial activity, carbon stock monitoring, and erosion measurements across contrasting climates and soil types. Such integrated experiments would clarify whether the thermal buffering observed here strengthens, weakens, or simply modifies the conditions controlling soil carbon persistence, biological activity, and ecosystem resilience.

## 5. Conclusion

This meta-analysis shows that biochar induces coordinated changes in soil thermal behaviour. Across 19 independent studies, biochar reduced thermal conductivity, thermal diffusivity, and volumetric heat capacity, while gravimetric heat capacity showed no significant overall response and remains too weakly represented to support firm inference. The strongest and most consistent evidence points to a structural mechanism: biochar reduces bulk density, modifies pore architecture, disrupts conductive pathways, and changes the volumetric composition of the soil matrix. Application rate, post-application bulk density, soil texture, moisture condition, feedstock group, and pyrolysis-related properties further modulated the magnitude of response, although these moderator effects should be interpreted as univariate associations.

These findings indicate that biochar-amended soils should not be treated as thermally unchanged systems. By modifying heat transfer, heat propagation, and volumetric heat storage, biochar may influence soil thermal buffering, soil moisture–temperature coupling, and temperature-sensitive biological and carbon-cycling processes. However, the field-scale implications remain insufficiently quantified, particularly for albedo, respiration, surface energy balance, and long-term carbon persistence. Future research should therefore integrate soil thermal measurements with hydrological, biological, and carbon-monitoring variables across contrasting climates, soils, and biochar types. Such integration will be essential for representing biochar effects in soil, land-surface, and carbon dioxide removal frameworks.

## Data, Materials, and Software Availability

The extracted data, harmonised analytical dataset, R analysis code, and figure-generation scripts supporting this study are publicly available in *Zenodo* at 10.5281/zenodo.20815506. The publications from which data were extracted are available through their original sources.

## Author Contributions

B.G. conceived and designed the study, conducted the literature search, screened studies, extracted and harmonised the data, performed the statistical analyses, generated the figures, and wrote the first draft. D.B. contributed to the statistical methodology, supervised the data analysis, and supported the interpretation of the meta-analytical results. K.W. contributed to the interpretation of soil thermal properties and biochar material characteristics. L.T. contributed to conceptual framing, resources, funding acquisition, and interpretation of soil–biochar interactions. O.M. contributed to the interpretation of biochar production, material properties, and carbon-removal implications. All authors critically reviewed and edited the manuscript and approved the final version.

## Acknowledgments

Open-access publication charges were supported by “The untapped potential of biochar for carbon mitigation”, funded by the Czech Ministry of Education, Youth and Sports, Czech Republic, under grant INTER-COST-LUC24045. B.G. and O.M. acknowledge support from the Horizon Europe C-SINK project, “Actions required to secure the large-scale deployment of the leading carbon dioxide removal (CDR) approaches to meet EU climate targets”, funded by the European Union under grant agreement number 101080377; https://doi.org/10.3030/101080377. The funders had no role in study design, data extraction, analysis, interpretation, or manuscript preparation.

## Primary classification

Physical Sciences — Environmental Sciences

## Secondary classification

Biological Sciences — Agricultural Sciences

## References

Azzi, E. S., Li, H., Cederlund, H., Karltun, E., & Sundberg, C. (2024). Modelling biochar long-term carbon storage in soil with harmonized analysis of decomposition data. Geoderma, 441, Article 116761. 10.1016/j.geoderma.2023.116761

Barbosa, T. A., Gomes Filho, R. R., Wisniewski, A., & Mašek, O. (2025). Biochar physical degradation: Long-term effects as soil amendments. Biomass and Bioenergy, 203, Article 108284. 10.1016/j.biombioe.2025.108284

Blanco-Canqui, H. (2017). Biochar and soil physical properties. Soil Science Society of America Journal, 81(4), 687–711. 10.2136/sssaj2017.01.0017

Bonan, G. (2019). Climate change and terrestrial ecosystem modeling. Cambridge University Press. 10.1017/9781107339217

Borenstein, M., Hedges, L. V., Higgins, J. P. T., & Rothstein, H. R. (2009). Introduction to meta-analysis. Wiley. 10.1002/9780470743386

Campbell, G. S., Jungbauer, J. D., Jr., Bidlake, W. R., & Hungerford, R. D. (1994). Predicting the effect of temperature on soil thermal conductivity. Soil Science, 158(5), 307–313. 10.1097/00010694-199411000-00001

Crombie, K., Mašek, O., Sohi, S. P., Brownsort, P., & Cross, A. (2013). The effect of pyrolysis conditions on biochar stability as determined by three methods. GCB Bioenergy, 5(2), 122–131. 10.1111/gcbb.12030

Dai, Y., Wei, N., Yuan, H., Zhang, S., Shangguan, W., Liu, S., Lu, X., & Xin, Y. (2019). Evaluation of soil thermal conductivity schemes for use in land surface modeling. Journal of Advances in Modeling Earth Systems, 11(11), 3454–3473. 10.1029/2019MS001723

de Vries, D. A. (1963). Thermal properties of soils. In Physics of plant environment. North-Holland.

Edeh, I. G., & Mašek, O. (2022). The role of biochar particle size and hydrophobicity in improving soil hydraulic properties. European Journal of Soil Science, 73(1), Article e13138. 10.1111/ejss.13138

Edeh, I. G., Mašek, O., & Buss, W. (2020). A meta-analysis on biochar’s effects on soil water properties: New insights and future research challenges. Science of the Total Environment, 714, 136857. 10.1016/j.scitotenv.2020.136857

Edeh, I. G., Mašek, O., & Fusseis, F. (2023). 4D structural changes and pore network model of biomass during pyrolysis. Scientific Reports, 13, Article 22863. 10.1038/s41598-023-49919-z

Fatichi, S., Or, D., Walko, R., Vereecken, H., Young, M. H., Ghezzehei, T. A., Hengl, T., Kollet, S., Agam, N., Avissar, R., & Maxwell, R. M. (2020). Soil structure is an important omission in Earth system models. Nature Communications, 11, 522. 10.1038/s41467-020-14411-z

Genesio, L., Miglietta, F., Baronti, S., & Vaccari, F. P. (2012). Surface albedo following biochar application in durum wheat. Environmental Research Letters, 7(1), Article 014025. 10.1088/1748-9326/7/1/014025

Gholamahmadi, B. (2026). Understanding feedstock-dependent biochar performance beyond static material descriptors. Biochar X, 2, Article e016. 10.48130/bchax-0026-0014

Gholamahmadi, B., & Kammann, C. (2026). Biochar for durable carbon removal: Soil erosion reduction as a key mechanism. Biomass Futures, 1, Article 100020. 10.1016/j.bmf.2026.100020

Gholamahmadi, B., Ferreira, C. S. S., Gonzalez-Pelayo, O., Bastos, A. C., & Verheijen, F. G. A. (2025a). Soil conservation benefits of biochar in Mediterranean vineyards: Enhancing the soil sponge function and mitigating water erosion. Biochar, 7, Article 106. 10.1007/s42773-025-00483-x

Gholamahmadi, B., Gonzalez-Pelayo, O., Isaka, S., Campos, I., Martins, M., Bastos, A. C., Jongen, M., & Verheijen, F. G. A. (2025b). The impact of biochar application on sponge function, water erosion, and vegetation cover in a Mediterranean vineyard soil. Journal of Environmental Management, 388, Article 125916. 10.1016/j.jenvman.2025.125916

Gholamahmadi, B., Jeffery, S., Gonzalez-Pelayo, O., Prats, S. A., Bastos, A. C., Keizer, J. J., & Verheijen, F. G. A. (2023). Biochar impacts on runoff and soil erosion by water: A systematic global-scale meta-analysis. Science of the Total Environment, 871, Article 161860. 10.1016/j.scitotenv.2023.161860

Ghorbani, M., & Amirahmadi, E. (2024). Insights into soil and biochar variations and their contribution to soil aggregate status – A meta-analysis. Soil & Tillage Research, 244, Article 106282. 10.1016/j.still.2024.106282

Hedges, L. V., Gurevitch, J., & Curtis, P. S. (1999). The meta-analysis of response ratios in experimental ecology. Ecology, 80, 1150–1156. https://doi.org/10.1890/0012-9658(1999)080[1150:TMAORR]20.CO;2

Hengl, T., Mendes de Jesus, J., Heuvelink, G. B. M., Ruiperez Gonzalez, M., Kilibarda, M., Blagotić, A., Shangguan, W., Wright, M. N., Geng, X., Bauer-Marschallinger, B., Guevara, M. A., Vargas, R., MacMillan, R. A., Batjes, N. H., Leenaars, J. G. B., Ribeiro, E., Wheeler, I., Mantel, S., & Kempen, B. (2017). SoilGrids250m: Global gridded soil information based on machine learning. PLOS ONE, 12(2), Article e0169748. 10.1371/journal.pone.0169748

Intergovernmental Panel on Climate Change. (2021). *Climate change 2021: The physical science basis: Contribution of Working Group I to the Sixth Assessment Report of the Intergovernmental Panel on Climate Change*. Cambridge University Press. 10.1017/9781009157896

Ivanova, N., Obaeed, G. L. O., Sulkarnaev, F., Buchkina, N., Gubin, A., & Yurtaev, A. (2023). Effect of biochar aging in agricultural soil on its wetting properties and surface structure. Biochar, 5, Article 75. 10.1007/s42773-023-00272-4

Khaledi, S., Delbari, M., Galavi, H., Bagheri, H., & Chari, M. M. (2023). Effects of biochar particle size, biochar application rate, and moisture content on thermal properties of an unsaturated sandy loam soil. Soil and Tillage Research, 226, Article 105579. 10.1016/j.still.2022.105579

Lawrence, D. M., Fisher, R. A., Koven, C. D., Oleson, K. W., Swenson, S. C., Bonan, G., Collier, N., Ghimire, B., van Kampenhout, L., Kennedy, D., Kluzek, E., Lawrence, P. J., Li, F., Li, H., Lombardozzi, D. L., Riley, W. J., Sacks, W. J., Shi, M., et al. (2019). The Community Land Model version 5: Description of new features, benchmarking, and impact of forcing uncertainty. Journal of Advances in Modeling Earth Systems, 11(12), 4245–4287. 10.1029/2018MS001583

Lehmann, J., & Joseph, S. (Eds.). (2015). Biochar for environmental management: Science, technology and implementation (2nd ed.). Routledge. 10.4324/9780203762264

Li, X., Wang, T., Chang, S. X., Jiang, X., & Song, Y. (2020). Biochar increases soil microbial biomass but has variable effects on microbial diversity: A meta-analysis. Science of the Total Environment, 749, Article 141593. 10.1016/j.scitotenv.2020.141593

Lu, Y., Lu, S., Horton, R., & Ren, T. (2014). An empirical model for estimating soil thermal conductivity from texture, water content, and bulk density. Soil Science Society of America Journal, 78(6), 1859–1868. 10.2136/sssaj2014.05.0218

Meyer, S., Bright, R. M., Fischer, D., Schulz, H., & Glaser, B. (2012). Albedo impact on the suitability of biochar systems to mitigate global warming. Environmental Science & Technology, 46(22), 12726–12734. 10.1021/es302302g

Ngambia, A., Mašek, O., & Erastova, V. (2024). Development of biochar molecular models with controlled porosity. Biomass and Bioenergy, 184, Article 107199. 10.1016/j.biombioe.2024.107199

Ochsner, T. E., Horton, R., & Ren, T. (2001). A new perspective on soil thermal properties. Soil Science Society of America Journal, 65(6), 1641–1647. 10.2136/sssaj2001.1641

Patwa, D., Bordoloi, U., Dubey, A. A., Ravi, K., Sekharan, S., & Kalita, P. (2022). Energy-efficient biochar production for thermal backfill applications. Science of the Total Environment, 833, Article 155253. 10.1016/j.scitotenv.2022.155253

Pokharel, P., Kwak, J.-H., Ok, Y. S., & Chang, S. X. (2020). Biochar improves soil properties and crop productivity: A meta-analysis. Agriculture, Ecosystems & Environment, 295, 106879. 10.1016/j.agee.2020.106879

Polak, M. P., & Morgan, D. (2025). Leveraging vision capabilities of multimodal large language models for automated data extraction from plots. arXiv. https://arxiv.org/abs/2503.12326

R Core Team. (2024). R: A language and environment for statistical computing. R Foundation for Statistical Computing. https://www.R-project.org/

Razzaghi, F., Obour, P. B., & Arthur, E. (2020). Does biochar improve soil water retention? A systematic review and meta-analysis. Geoderma, 361, Article 114055. 10.1016/j.geoderma.2019.114055

Schmidt, H.-P., Kammann, C., Hagemann, N., Leifeld, J., Bucheli, T. D., Sánchez Monedero, M. A., & Cayuela, M. L. (2021). Biochar in agriculture: A systematic review of 26 global meta-analyses. GCB Bioenergy, 13(11), 1708–1730. 10.1111/gcbb.12889

Shi, G., Sun, W., Shangguan, W., Wei, Z., Yuan, H., Li, L., Sun, X., Zhang, Y., Liang, H., Li, D., Huang, F., Li, Q., & Dai, Y. (2025). A China dataset of soil properties for land surface modelling (version 2, CSDLv2). Earth System Science Data, 17, 517–543. 10.5194/essd-17-517-2025

Tong, B., Gao, Z., Horton, R., Li, Y., & Wang, L. (2016). An empirical model for estimating soil thermal conductivity from soil water content and porosity. Journal of Hydrometeorology, 17(2), 601–613. 10.1175/JHM-D-15-0119.1

Usowicz, B., Lipiec, J., Łukowski, M., Bis, Z., Usowicz, J., & Latawiec, A. E. (2020). Impact of biochar addition on soil thermal properties: Modelling approach. Geoderma, 376, Article 114574. 10.1016/j.geoderma.2020.114574

Usowicz, B., Lipiec, J., Łukowski, M., Marczewski, W., & Usowicz, J. (2016). The effect of biochar application on thermal properties and albedo of loess soil under grassland and fallow. *Soil and Tillage Research, 164*, 45–51. 10.1016/j.still.2016.03.009

Verheijen, F. G. A., Jeffery, S., van der Velde, M., Penížek, V., Beland, M., Bastos, A. C., & Keizer, J. J. (2013). Reductions in soil surface albedo as a function of biochar application rate: Implications for global radiative forcing. Environmental Research Letters, 8(4), Article 044008. 10.1088/1748-9326/8/4/044008

Viechtbauer, W. (2010). Conducting meta-analyses in R with the metafor package. Journal of Statistical Software, 36(3), 1–48. 10.18637/jss.v036.i03

Wang, D., Li, C., Parikh, S. J., & Scow, K. M. (2019). Impact of biochar on water retention of two agricultural soils: A multi-scale analysis. Geoderma, 340, 185– 191. 10.1016/j.geoderma.2019.01.012

Wang, M., Yu, X., Weng, X., Zeng, X., Li, M., & Sui, X. (2023). Meta-analysis of the effects of biochar application on the diversity of soil bacteria and fungi. Microorganisms, 11(3), Article 641. 10.3390/microorganisms11030641

Weber, K., & Quicker, P. (2018). Properties of biochar. Fuel, 217, 240–261. 10.1016/j.fuel.2017.12.054

Weng, Z. H., & Cowie, A. L. (2025). Estimates vary but credible evidence points to gigaton-scale climate change mitigation potential of biochar. Communications Earth & Environment, 6, Article 259. 10.1038/s43247-025-02228-x

Woolf, D., Amonette, J. E., Street-Perrott, F. A., Lehmann, J., & Joseph, S. (2010). Sustainable biochar to mitigate global climate change. *Nature Communications, 1*, Article 56. 10.1038/ncomms1053

Woolf, D., Lehmann, J., Ogle, S., Kishimoto-Mo, A. W., McConkey, B., & Baldock, J. (2021). Greenhouse gas inventory model for biochar additions to soil. Environmental Science & Technology, 55(21), 14795–14805. 10.1021/acs.est.1c02425

Xie, X., Lu, Y., Ren, T., & Horton, R. (2020). Thermal conductivity of mineral soils relates linearly to air-filled porosity. *Soil Science Society of America Journal, 84*(1), 53–56. 10.1002/saj2.20016

Xiong, J., Yu, R., Islam, E., Zhu, F., Zha, J., & Sohail, M. I. (2020). Effect of biochar on soil temperature under high soil surface temperature in coal mined arid and semiarid regions. Sustainability, 12(19), Article 8238. 10.3390/su12198238

Xu, W., Xu, H., Delgado-Baquerizo, M., Gundale, M. J., Zou, X., & Ruan, H. (2023). Global meta-analysis reveals positive effects of biochar on soil microbial diversity. Geoderma, 436, Article 116528. 10.1016/j.geoderma.2023.116528

Zhang, Q., Wang, Y., Wu, Y., Wang, X., Du, Z., Liu, X., & Song, J. (2013). Effects of biochar amendment on soil thermal conductivity, reflectance, and temperature. Soil Science Society of America Journal, 77(5), 1478–1487. 10.2136/sssaj2012.0180

Zhao, B. W., Zhao, Y., Liu, H., Li, Y. Q., Duan, K. X., & Zhang, X. (2021). Effect of wheat straw biochar on thermophysical properties of loessial soil. Nature Environment and Pollution Technology, 20(3), 1033–1039. 10.46488/NEPT.2021.v20i03.010

Zhao, J., Ren, T., Zhang, Q., Du, Z., & Wang, Y. (2016). Effects of biochar amendment on soil thermal properties in the North China Plain. *Soil Science Society of America Journal, 80*(5), 1157–1166. 10.2136/sssaj2016.01.0020

Zhao, L., Cao, X., Mašek, O., & Zimmerman, A. (2013). Heterogeneity of biochar properties as a function of feedstock sources and production temperatures. Journal of Hazardous Materials, 256–257, 1–9. 10.1016/j.jhazmat.2013.04.015

