## Supplementary Information for "Biochar reduces soil thermal conductivity, diffusivity and volumetric heat storage: A global meta-analysis"

**Contents:**

Figure S1. PRISMA flow diagram of study identification, screening, eligibility assessment, and inclusion for the meta-analysis of biochar effects on soil thermal properties.

Figure S2. Global distribution of studies included in the meta-analysis. Pins indicate unique study locations for the 19 independent studies included in the quantitative synthesis. The uneven distribution of study locations reflects the available experimental evidence base.

Figure S3. Small-study-effect and funnel-asymmetry diagnostics

Supplementary Methods S1. PRISMA-based study identification and screening

Supplementary Methods S2. Random-effects structure comparison

Supplementary Methods S3. Moderator screening

Supplementary Methods S4. Leave-one-study-out sensitivity analysis

Supplementary Methods S5. Small-study effects and funnel asymmetry diagnostics

Table S1. PRISMA flow summary

Table S2. Comparison of alternative random-effects structures for meta-analytic models of biochar effects on soil thermal properties. M1 includes an effect-size-level heterogeneity component, M2 includes study-level heterogeneity only, and M3 includes both study-level and within-study effect-size heterogeneity. Values shown are from restricted maximum-likelihood model fits. Lower AIC, AICc, and BIC values indicate better model fit.

Table S3. Robust moderator screening results for biochar effects on soil thermal properties

Table S4. Coefficients for selected continuous moderators of biochar effects on soil thermal properties

Table S5. Categorical moderator level estimates for selected biochar, soil, and climate moderators

Table S6. Pairwise co-variation among biochar-induced changes in soil thermal properties

Table S7. Summary of leave-one-study-out sensitivity analyses

Table S8. Small-study effects and funnel asymmetry diagnostics for biochar effects on soil thermal properties. Egger-type models used the standard error of the log response ratio as a moderator within the three-level random-effects framework. Precision-based models were fitted as sensitivity diagnostics. Significant slopes indicate potential small-study effects or funnel asymmetry, but should not be interpreted as definitive evidence of publication bias.

PRISMA flow diagram of study identification and selection

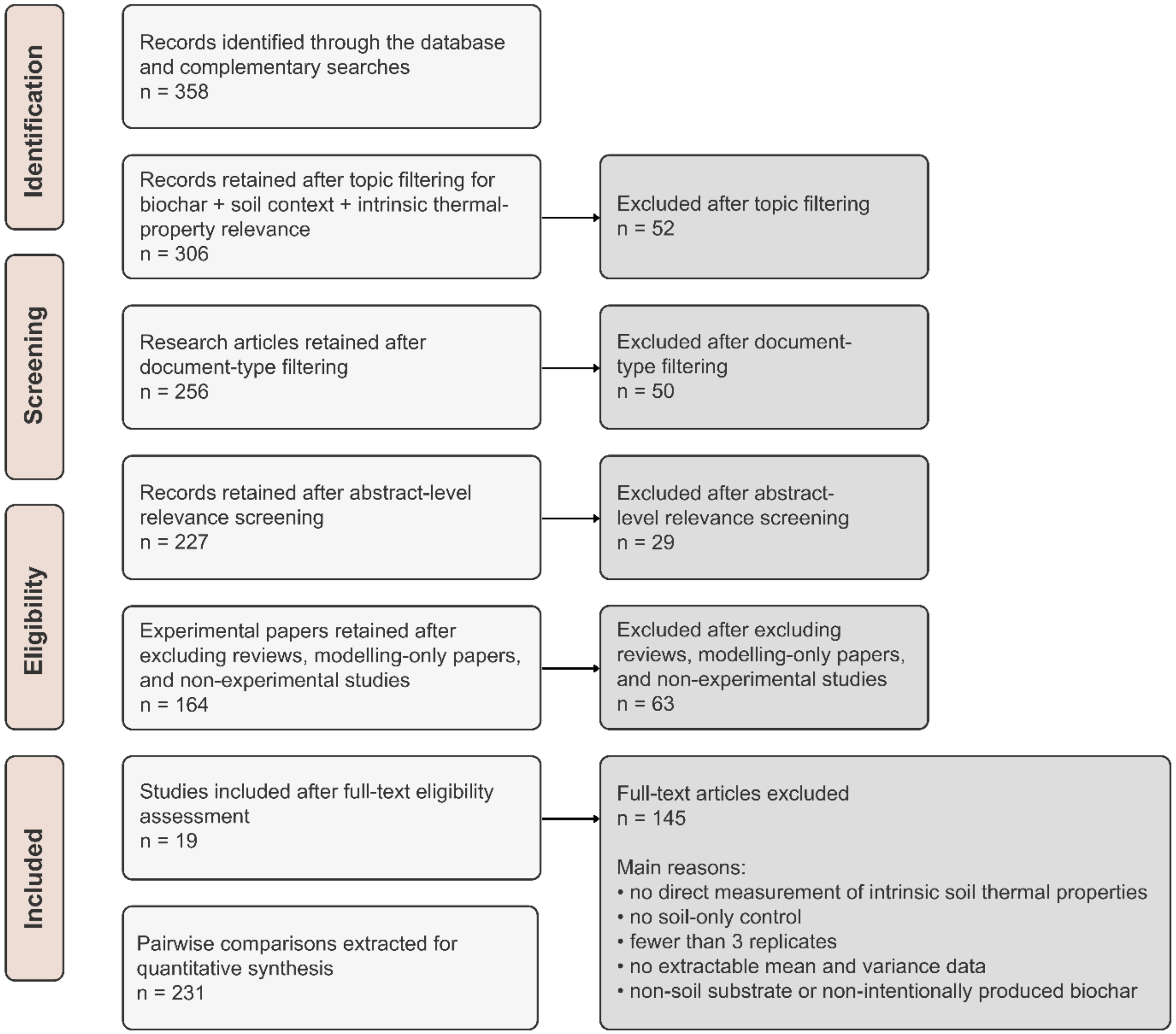

Figure S1. PRISMA flow diagram of study identification, screening, eligibility assessment, and inclusion in the meta-analysis of biochar effects on soil thermal properties.

### Supplementary Methods S1. PRISMA-based study identification and screening

The systematic review and study selection process followed PRISMA 2020 guidelines. The literature search initially identified 358 records from database searches and complementary screening. Database searches were performed in Web of Science Core Collection and Scopus, and Google Scholar was used as a complementary source by screening the first 200 results sorted by relevance, with screening stopped when no additional eligible records were identified. Records were first filtered for relevance to the combined topic of biochar, soil context, and intrinsic soil thermal properties, while excluding engineering- or materials-focused studies without a soil experimental context. This first screening step retained 306 records. The record set was then restricted to peer-reviewed research articles, reducing the number of records to 256. Abstract-level screening was subsequently conducted to identify studies directly relevant to biochar effects on soil thermal behaviour, retaining 227 records. Reviews, modelling-only papers, and non-experimental studies were then excluded, leaving 164 experimental papers for full-text eligibility assessment.

Full-text assessment was guided by predefined inclusion and exclusion criteria. Studies were included if they experimentally applied intentionally produced biochar to soil under laboratory, pot, column, or field conditions; measured at least one intrinsic soil thermal property, including thermal conductivity, thermal diffusivity, volumetric heat capacity, gravimetric heat capacity, or thermal resistivity; included a soil-only control without biochar amendment; reported mean values for both control and biochar-amended treatments; included at least three independent replicates per treatment; and reported a measure of variability, such as standard deviation, standard error, or confidence interval, or provided sufficient information to derive standard deviations without statistical imputation.

Studies were excluded if they reported soil or surface temperature time series without direct measurement of intrinsic thermal properties; investigated historical, technogenic, or naturally occurring charcoal rather than intentionally produced biochar; used non-soil substrates such as engineered media, hydroponic mixes, or construction materials; lacked a soil-only control treatment; used fewer than three replicates; did not report variability measures and did not allow variance estimation; or provided incomplete outcomes, such as missing control or treatment means. No statistical imputation was applied for missing sample size or variance.

Of the 164 experimental papers assessed at full text, 145 were excluded because they did not meet one or more eligibility criteria. The final quantitative synthesis, therefore included 19 independent studies, from which 231 unique pairwise control–biochar comparisons were extracted. Because several comparisons reported more than one thermal property, these comparisons yielded 529 property-specific effect sizes after excluding six volumetric heat capacity observations with non-positive sampling variance. The final property-specific dataset included 231 effect sizes for thermal conductivity, 118 for thermal diffusivity, 122 for volumetric heat capacity, and 58 for gravimetric heat capacity.

Table S1. PRISMA flow summary

| Screening stage | Records/studies retained | Records/studies excluded at stage |
|---|---|---|
| Records identified through database and complementary searches | 358 | — |
| Records retained after topic filtering for biochar + soil context + intrinsic thermal-property relevance | 306 | 52 |
| Research articles retained after document-type filtering | 256 | 50 |
| Records retained after abstract-level relevance screening | 227 | 29 |
| Experimental papers retained after excluding reviews, modelling-only papers, and non-experimental studies | 164 | 63 |
| Studies included after full-text eligibility assessment | 19 | 145 |
| Unique pairwise control–biochar comparisons extracted | 231 | — |
| Property-specific effect sizes retained after variance screening | 529 | 6 |

**Global evidence base for biochar effects on soil thermal properties**

Interactive map of the 19 independent studies included in the meta-analysis. Click a pin to view the study, year, DOI and measured location.

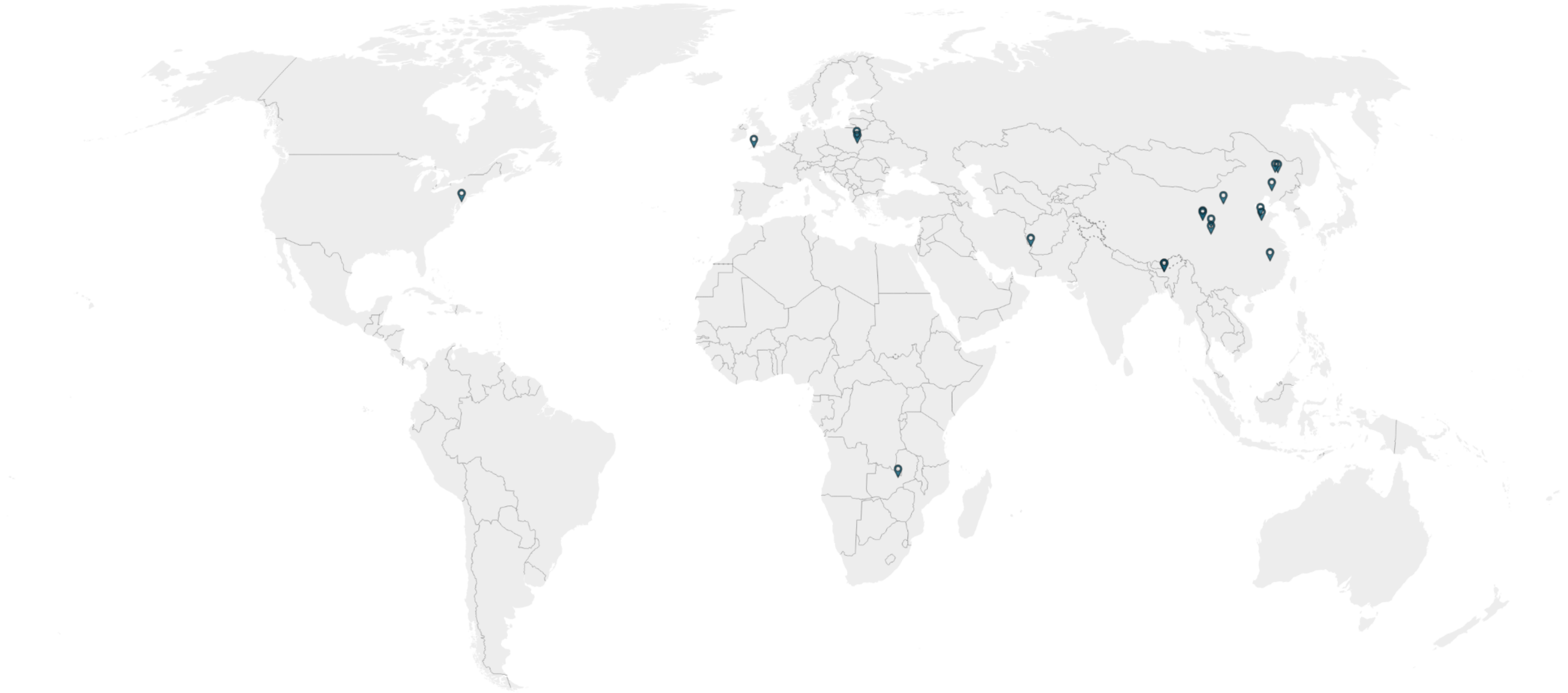

Source: Gholamahmadi, Beillouin, Weber, Trakal, Mašek "Biochar reduces soil thermal conductivity, diffusivity and volumetric heat storage: A global meta-analysis" • Created with Datawrapper

Figure S2. Global distribution of studies included in the meta-analysis. Pins indicate unique study locations for the 19 independent studies included in the quantitative synthesis. An interactive version provides study-level metadata, including study citation, DOI and geographic coordinates. Link: https://datawrapper.dwcdn.net/qs20l/1/

### Supplementary Methods S2. Random-effects structure comparison

Three alternative random-effects structures were compared for each thermal property: (i) a conventional model with effect-size-level heterogeneity, (ii) a model containing study-level heterogeneity only, and (iii) a three-level model containing both study-level and within-study effect-size heterogeneity. Across all four thermal properties, the three-level model consistently produced the lowest AIC, AICc, and BIC values under restricted maximum likelihood. The same model ranking was obtained using maximum likelihood as a sensitivity check, confirming that the preferred random-effects structure was robust to estimator choice. The three-level structure was therefore retained for the primary analyses.

Table S2. Comparison of alternative random-effects structures for meta-analytic models of biochar effects on soil thermal properties. M1 includes an effect-size-level heterogeneity component, M2 includes study-level heterogeneity only, and M3 includes both study-level and within-study effect-size heterogeneity. Values shown are from restricted maximum-likelihood model fits. Lower AIC, AICc, and BIC values indicate better model fit.

| thermal_property | estimator | model | n_effect_sizes | n_studies | estimate_lnRR | logLik | AIC | AICc | BIC |
|---|---|---|---|---|---|---|---|---|---|
| k | REML | M1: effect-size-level heterogeneity | 231 | 19 | -0.203 | -1.2 | 6.3 | 6.4 | 13.2 |
| k | REML | M2: study-level heterogeneity only | 231 | 19 | -0.181 | -1697.6 | 3399.2 | 3399.2 | 3406.1 |
| k | REML | M3: effect sizes nested within study | 231 | 19 | -0.194 | 59.0 | -112.0 | -111.9 | -101.7 |
| α | REML | M1: effect-size-level heterogeneity | 118 | 12 | -0.106 | 98.4 | -192.9 | -192.8 | -187.4 |
| α | REML | M2: study-level heterogeneity only | 118 | 12 | -0.111 | -520.6 | 1045.3 | 1045.4 | 1050.8 |

| | | | | | | | | | |
|---|---|---|---|---|---|---|---|---|---|
| α | REML | M3: effect sizes nested within study | 118 | 12 | -0.116 | 106.6 | -207.2 | -207.0 | -199.0 |
| Cv | REML | M1: effect-size-level heterogeneity | 122 | 13 | -0.087 | 106.2 | -208.4 | -208.3 | -202.9 |
| Cv | REML | M2: study-level heterogeneity only | 122 | 13 | -0.083 | -1075.0 | 2154.0 | 2154.1 | 2159.6 |
| Cv | REML | M3: effect sizes nested within study | 122 | 13 | -0.087 | 125.0 | -244.0 | -243.8 | -235.6 |
| Cp | REML | M1: effect-size-level heterogeneity | 58 | 6 | 0.015 | 30.6 | -57.2 | -56.9 | -53.1 |
| Cp | REML | M2: study-level heterogeneity only | 58 | 6 | 0.035 | -176.0 | 356.0 | 356.2 | 360.1 |
| Cp | REML | M3: effect sizes nested within study | 58 | 6 | 0.032 | 61.7 | -117.4 | -116.9 | -111.3 |

Table S3. Moderator screening results for biochar effects on soil thermal properties. Moderator effects were tested using univariate meta-regression models within the three-level random-effects framework. Continuous moderators were analysed as numeric predictors. Categorical moderators were retained only when at least two categories had sufficient replication, defined as at least five effect sizes and at least two independent studies per category. Variables with strong study-specific structure were treated as exploratory and not used for main-text interpretation.

| thermal_property | moderator | moderator_type | n_effect_sizes | n_studies | n_levels | QM | QM_p | AIC | BIC |
|---|---|---|---|---|---|---|---|---|---|
| Cp | delta_bulk_density_percent | continuous | 54 | 5 | NA | 52.46 | <0.001 | -137.1 | -129.3 |
| Cp | application_rate_t_ha | continuous | 27 | 4 | NA | 34.19 | <0.001 | -52.1 | -47.2 |
| Cp | post_application_soil_bulk_density_g_cm3 | continuous | 54 | 5 | NA | 11.87 | <0.001 | -114.0 | -106.2 |
| Cp | soil_porosity_percent | continuous | 27 | 4 | NA | 11.46 | <0.001 | -40.6 | -35.7 |
| Cp | usda_sand_percent | continuous | 49 | 5 | NA | 8.18 | 0.004 | -119.5 | -112.1 |
| Cp | usda_silt_percent | continuous | 58 | 6 | NA | 6.29 | 0.012 | -120.1 | -112.0 |
| Cp | usda_clay_percent | continuous | 58 | 6 | NA | 3.38 | 0.066 | -118.6 | -110.5 |
| Cp | application_rate_w_w_percent | continuous | 58 | 6 | NA | 2.82 | 0.093 | -114.9 | -106.8 |
| Cp | duration_days | continuous | 58 | 6 | NA | 1.21 | 0.271 | -116.4 | -108.3 |
| Cp | pyrolysis_temperature_group | categorical | 55 | 6 | 2 | 0.49 | 0.483 | -111.7 | -103.8 |
| Cp | measurement_temp_c | continuous | 52 | 5 | NA | 0.48 | 0.490 | -107.5 | -99.9 |
| Cp | heating_rate_c_per_min | continuous | 44 | 4 | NA | 0.39 | 0.533 | -92.7 | -85.7 |
| Cp | biochar_c_percent | continuous | 52 | 5 | NA | 0.29 | 0.589 | -100.4 | -92.8 |
| Cp | pyrolysis_temperature_c | continuous | 44 | 4 | NA | 0.24 | 0.622 | -90.0 | -83.1 |
| Cp | soil_bulk_density_g_cm3 | continuous | 58 | 6 | NA | 0.20 | 0.653 | -114.8 | -106.7 |
| Cp | pyrolysis_duration_h | continuous | 44 | 4 | NA | 0.16 | 0.692 | -93.2 | -86.3 |
| Cp | biochar_p_h | continuous | 27 | 4 | NA | 0.07 | 0.789 | -30.7 | -25.8 |
| Cp | soil_p_h | continuous | 27 | 4 | NA | 0.06 | 0.811 | -37.0 | -32.1 |
| Cv | soil_texture_group | categorical | 97 | 9 | 3 | 23.56 | <0.001 | -228.8 | -216.1 |
| Cv | post_application_soil_bulk_density_g_cm3 | continuous | 102 | 11 | NA | 18.77 | <0.001 | -225.2 | -214.7 |

| | | | | | | | | | |
|---|---|---|---|---|---|---|---|---|---|
| Cv | biochar_particle_size_group | categorical | 65 | 9 | 2 | 16.65 | <0.001 | -103.8 | -95.3 |
| Cv | feedstock_group | categorical | 110 | 12 | 3 | 17.52 | <0.001 | -209.7 | -196.4 |
| Cv | soil_moisture_percent | continuous | 69 | 6 | NA | 13.11 | <0.001 | -180.2 | -171.4 |
| Cv | application_rate_w_w_percent | continuous | 122 | 13 | NA | 10.73 | 0.001 | -249.2 | -238.1 |
| Cv | measurement_depth_down_to_cm | continuous | 26 | 4 | NA | 9.41 | 0.002 | -84.6 | -79.9 |
| Cv | delta_bulk_density_percent | continuous | 98 | 10 | NA | 9.25 | 0.002 | -218.4 | -208.2 |
| Cv | usda_clay_percent | continuous | 122 | 13 | NA | 8.66 | 0.003 | -248.3 | -237.1 |
| Cv | surface_area_m2_g | continuous | 94 | 8 | NA | 8.18 | 0.004 | -212.2 | -202.1 |
| Cv | climate_zone_by_lat_and_long | categorical | 122 | 13 | 2 | 6.33 | 0.012 | -245.9 | -234.8 |
| Cv | pyrolysis_duration_h | continuous | 95 | 8 | NA | 6.00 | 0.014 | -211.6 | -201.4 |
| Cv | soil_bulk_density_g_cm3 | continuous | 102 | 11 | NA | 5.62 | 0.018 | -225.4 | -215.0 |
| Cv | usda_sand_percent | continuous | 113 | 12 | NA | 4.84 | 0.028 | -223.7 | -212.8 |
| Cv | soc_percent | continuous | 40 | 5 | NA | 3.77 | 0.052 | -85.4 | -78.9 |
| Cv | soil_p_h | continuous | 67 | 8 | NA | 3.44 | 0.064 | -126.8 | -118.1 |
| Cv | soil_porosity_percent | continuous | 61 | 9 | NA | 2.45 | 0.117 | -133.4 | -125.1 |
| Cv | heating_rate_c_per_min | continuous | 95 | 8 | NA | 2.10 | 0.147 | -208.3 | -198.1 |
| Cv | duration_days | continuous | 122 | 13 | NA | 1.32 | 0.251 | -241.9 | -230.8 |
| Cv | measurement_temp_c | continuous | 93 | 10 | NA | 1.10 | 0.293 | -205.1 | -195.0 |
| Cv | pyrolysis_temperature_c | continuous | 101 | 10 | NA | 0.73 | 0.393 | -216.4 | -206.0 |
| Cv | biochar_c_percent | continuous | 94 | 10 | NA | 0.72 | 0.395 | -191.2 | -181.1 |
| Cv | application_rate_t_ha | continuous | 35 | 6 | NA | 0.54 | 0.463 | -58.2 | -52.2 |

| | | | | | | | | | |
|---|---|---|---|---|---|---|---|---|---|
| Cv | usda_silt_percent | continuous | 122 | 13 | NA | 0.53 | 0.467 | -241.5 | -230.3 |
| Cv | pyrolysis_temperature_group | categorical | 122 | 13 | 3 | 0.99 | 0.608 | -235.3 | -221.4 |
| Cv | biochar_p_h | continuous | 77 | 9 | NA | 0.26 | 0.610 | -148.6 | -139.3 |
| α | delta_bulk_density_percent | continuous | 104 | 10 | NA | 27.62 | <0.001 | -213.2 | -202.7 |
| α | application_rate_w_w_percent | continuous | 118 | 12 | NA | 15.70 | <0.001 | -216.4 | -205.4 |
| α | post_application_soil_bulk_density_g_cm3 | continuous | 108 | 11 | NA | 10.04 | 0.002 | -196.2 | -185.5 |
| α | measurement_temp_c | continuous | 99 | 10 | NA | 8.37 | 0.004 | -184.5 | -174.2 |
| α | surface_area_m2_g | continuous | 84 | 7 | NA | 7.72 | 0.005 | -202.8 | -193.1 |
| α | soil_bulk_density_g_cm3 | continuous | 108 | 11 | NA | 6.79 | 0.009 | -202.0 | -191.3 |
| α | soc_percent | continuous | 46 | 5 | NA | 6.38 | 0.012 | -58.5 | -51.4 |
| α | pyrolysis_temperature_c | continuous | 91 | 9 | NA | 6.29 | 0.012 | -169.0 | -159.1 |
| α | pyrolysis_temperature_group | categorical | 118 | 12 | 3 | 7.91 | 0.019 | -204.7 | -191.0 |
| α | biochar_c_percent | continuous | 90 | 9 | NA | 4.74 | 0.029 | -193.8 | -183.9 |
| α | usda_clay_percent | continuous | 118 | 12 | NA | 2.49 | 0.114 | -205.5 | -194.5 |
| α | soil_p_h | continuous | 73 | 8 | NA | 2.31 | 0.128 | -123.9 | -114.8 |
| α | application_rate_t_ha | continuous | 41 | 6 | NA | 2.26 | 0.133 | -62.2 | -55.6 |
| α | soil_texture_group | categorical | 88 | 8 | 2 | 1.81 | 0.179 | -199.0 | -189.2 |
| α | pyrolysis_duration_h | continuous | 85 | 7 | NA | 1.73 | 0.188 | -155.8 | -146.2 |
| α | soil_porosity_percent | continuous | 67 | 9 | NA | 1.50 | 0.220 | -143.6 | -134.9 |
| α | usda_silt_percent | continuous | 118 | 12 | NA | 1.03 | 0.310 | -205.0 | -194.0 |
| α | biochar_p_h | continuous | 73 | 8 | NA | 0.83 | 0.361 | -163.6 | -154.5 |

| | | | | | | | | | |
|---|---|---|---|---|---|---|---|---|---|
| α | duration_days | continuous | 118 | 12 | NA | 0.76 | 0.383 | -204.4 | -193.4 |
| α | feedstock_group | categorical | 102 | 11 | 2 | 0.49 | 0.483 | -156.9 | -146.5 |
| α | usda_sand_percent | continuous | 109 | 11 | NA | 0.34 | 0.560 | -184.6 | -173.9 |
| α | soil_moisture_percent | continuous | 69 | 6 | NA | 0.24 | 0.622 | -110.9 | -102.0 |
| α | biochar_particle_size_group | categorical | 61 | 8 | 2 | 0.13 | 0.719 | -70.3 | -62.0 |
| α | heating_rate_c_per_min | continuous | 85 | 7 | NA | 0.02 | 0.882 | -154.5 | -144.8 |
| α | climate_zone_by_lat_and_long | categorical | 118 | 12 | 2 | 0.00 | 0.958 | -204.2 | -193.2 |
| α | measurement_depth_down_to_cm | continuous | 26 | 4 | NA | 0.00 | 0.962 | -62.4 | -57.6 |
| k | application_rate_w_w_percent | continuous | 231 | 19 | NA | 254.43 | <0.001 | -335.8 | -322.1 |
| k | post_application_soil_bulk_density_g_cm3 | continuous | 161 | 14 | NA | 29.08 | <0.001 | -265.4 | -253.1 |
| k | delta_bulk_density_percent | continuous | 157 | 13 | NA | 24.13 | <0.001 | -264.8 | -252.6 |
| k | measurement_temp_c | continuous | 158 | 14 | NA | 11.70 | <0.001 | -248.2 | -236.0 |
| k | feedstock_group | categorical | 219 | 18 | 3 | 10.66 | 0.005 | -90.0 | -73.1 |
| k | soc_percent | continuous | 133 | 7 | NA | 6.48 | 0.011 | -4.3 | 7.2 |
| k | soil_texture_group | categorical | 209 | 15 | 4 | 9.49 | 0.023 | -123.1 | -103.2 |
| k | climate_zone_by_lat_and_long | categorical | 189 | 18 | 2 | 4.37 | 0.037 | -258.1 | -245.1 |
| k | soil_porosity_percent | continuous | 122 | 13 | NA | 4.26 | 0.039 | -251.8 | -240.6 |
| k | pyrolysis_duration_h | continuous | 184 | 11 | NA | 3.89 | 0.049 | -65.0 | -52.2 |
| k | measurement_depth_down_to_cm | continuous | 78 | 8 | NA | 3.50 | 0.062 | 52.4 | 61.7 |
| k | biochar_particle_size_group | categorical | 88 | 11 | 3 | 5.40 | 0.067 | -56.7 | -44.5 |
| k | heating_rate_c_per_min | continuous | 184 | 11 | NA | 2.85 | 0.091 | -64.2 | -51.4 |

| | | | | | | | | | |
|---|---|---|---|---|---|---|---|---|---|
| k | surface_area_m2_g | continuous | 139 | 9 | NA | 2.70 | 0.100 | -239.3 | -227.7 |
| k | usda_sand_percent | continuous | 222 | 18 | NA | 2.37 | 0.124 | -98.8 | -85.2 |
| k | ash_content_percent | continuous | 53 | 6 | NA | 2.14 | 0.144 | -51.8 | -44.1 |
| k | usda_clay_percent | continuous | 231 | 19 | NA | 1.25 | 0.264 | -110.6 | -96.9 |
| k | water_filled_pore_space_percent | continuous | 59 | 4 | NA | 1.20 | 0.274 | -127.8 | -119.7 |
| k | soil_moisture_percent | continuous | 119 | 10 | NA | 0.34 | 0.562 | -176.7 | -165.7 |
| k | pyrolysis_temperature_group | categorical | 231 | 19 | 3 | 1.12 | 0.570 | -105.7 | -88.6 |
| k | application_rate_t_ha | continuous | 51 | 9 | NA | 0.31 | 0.578 | -70.1 | -62.5 |
| k | biochar_p_h | continuous | 184 | 14 | NA | 0.30 | 0.586 | -81.1 | -68.2 |
| k | biochar_c_percent | continuous | 201 | 15 | NA | 0.15 | 0.696 | -112.2 | -99.0 |
| k | pyrolysis_temperature_c | continuous | 196 | 14 | NA | 0.11 | 0.741 | -77.4 | -64.4 |
| k | usda_silt_percent | continuous | 231 | 19 | NA | 0.06 | 0.800 | -110.2 | -96.4 |
| k | soil_bulk_density_g_cm3 | continuous | 209 | 16 | NA | 0.02 | 0.885 | -105.5 | -92.2 |
| k | soil_p_h | continuous | 166 | 11 | NA | 0.02 | 0.896 | -29.7 | -17.3 |
| k | duration_days | continuous | 231 | 19 | NA | 0.01 | 0.906 | -109.7 | -96.0 |

**Supplementary Methods S3. Moderator screening**

Moderator screening was conducted using univariate meta-regression models fitted within the same three-level random-effects structure as the main analysis. Candidate moderators included soil, biochar, experimental, and measurement-related variables. Continuous moderators were analysed as numeric predictors. Categorical moderators were retained for robust interpretation only when at least two categories had sufficient replication, defined as at least five effect sizes and at least two independent studies per category.

Variables with many study-specific levels, including individual experiment-type labels, incorporation method, thermal measurement method, and vegetation type, were screened exploratorily but were not used as primary evidence in the main text because they were closely confounded with study identity. Moderator results were therefore interpreted based on statistical support, replication, and mechanistic plausibility. Because these analyses were univariate and many moderator variables were incompletely reported or partially confounded, they should be interpreted as evidence for plausible associations rather than as definitive causal attribution.

Table S4. Coefficients for selected continuous moderators of biochar effects on soil thermal properties. Slopes are reported on the log response ratio scale and as approximate percentage change per one-unit increase in the moderator. Positive slopes indicate that the biochar effect becomes less negative or more positive as the moderator increases, whereas negative slopes indicate stronger reductions relative to the unamended control.

| thermal_property | moderator | n_effect_sizes | n_studies | intercept_lnrr | slope_lnrr | slope_ci_lb | slope_ci_ub | slope_p | slope_percent_per_unit | slope_ci_lb_percent | slope_ci_ub_percent |
|---|---|---|---|---|---|---|---|---|---|---|---|
| k | application_rate_w_w_percent | 231 | 19 | -0.120 | -0.013 | -0.015 | -0.012 | <0.001 | -1.323 | -1.485 | -1.162 |
| k | post_application_soil_bulk_density_g_cm3 | 161 | 14 | -0.644 | 0.392 | 0.250 | 0.535 | <0.001 | 48.041 | 28.368 | 70.729 |
| k | delta_bulk_density_percent | 157 | 13 | -0.085 | 0.007 | 0.004 | 0.009 | <0.001 | 0.656 | 0.394 | 0.919 |
| k | measurement_temp_c | 158 | 14 | 0.035 | -0.008 | -0.013 | -0.003 | <0.001 | -0.812 | -1.275 | -0.348 |
| k | soc_percent | 133 | 7 | -0.486 | 0.227 | 0.052 | 0.401 | 0.011 | 25.423 | 5.352 | 49.318 |
| k | soil_porosity_percent | 122 | 13 | 0.137 | -0.005 | -0.010 | 0.000 | 0.039 | -0.522 | -1.014 | -0.026 |
| k | pyrolysis_duration_h | 184 | 11 | -0.278 | 0.015 | 0.000 | 0.029 | 0.049 | 1.480 | 0.009 | 2.972 |

| | | | | | | | | | | | |
|---|---|---|---|---|---|---|---|---|---|---|---|
| α | delta_bulk_density_percent | 104 | 10 | -0.048 | 0.008 | 0.005 | 0.011 | <0.001 | 0.789 | 0.494 | 1.084 |
| α | application_rate_w_w_percent | 118 | 12 | -0.084 | -0.011 | -0.016 | -0.006 | <0.001 | -1.089 | -1.623 | -0.552 |
| α | post_application_soil_bulk_density_g_cm3 | 108 | 11 | -0.423 | 0.241 | 0.092 | 0.390 | 0.002 | 27.258 | 9.631 | 47.720 |
| α | measurement_temp_c | 99 | 10 | 0.021 | -0.006 | -0.009 | -0.002 | 0.004 | -0.550 | -0.921 | -0.178 |
| α | surface_area_m2_g | 84 | 7 | -0.106 | 0.000 | 0.000 | 0.001 | 0.005 | 0.034 | 0.010 | 0.058 |
| α | soil_bulk_density_g_cm3 | 108 | 11 | 0.087 | -0.145 | -0.254 | -0.036 | 0.009 | -13.513 | -22.457 | -3.538 |
| α | soc_percent | 46 | 5 | -0.300 | 0.158 | 0.035 | 0.281 | 0.012 | 17.115 | 3.605 | 32.387 |
| α | pyrolysis_temperature_c | 91 | 9 | -0.206 | 0.000 | 0.000 | 0.000 | 0.012 | 0.022 | 0.005 | 0.038 |
| α | biochar_c_percent | 90 | 9 | -0.229 | 0.002 | 0.000 | 0.003 | 0.029 | 0.184 | 0.018 | 0.349 |
| Cv | post_application_soil_bulk_density_g_cm3 | 102 | 11 | -0.457 | 0.306 | 0.168 | 0.445 | <0.001 | 35.858 | 18.270 | 56.061 |
| Cv | soil_moisture_percent | 69 | 6 | -0.157 | 0.002 | 0.001 | 0.003 | <0.001 | 0.181 | 0.083 | 0.279 |
| Cv | application_rate_w_w_percent | 122 | 13 | -0.058 | -0.009 | -0.014 | -0.003 | 0.001 | -0.865 | -1.379 | -0.349 |
| Cv | measurement_depth_down_to_cm | 26 | 4 | 0.022 | -0.013 | -0.021 | -0.005 | 0.002 | -1.299 | -2.119 | -0.471 |
| Cv | delta_bulk_density_percent | 98 | 10 | -0.030 | 0.005 | 0.002 | 0.008 | 0.002 | 0.463 | 0.164 | 0.762 |
| Cv | usda_clay_percent | 122 | 13 | -0.044 | -0.003 | -0.005 | -0.001 | 0.003 | -0.277 | -0.462 | -0.093 |
| Cv | surface_area_m2_g | 94 | 8 | -0.120 | 0.000 | 0.000 | 0.001 | 0.004 | 0.038 | 0.012 | 0.064 |
| Cv | pyrolysis_duration_h | 95 | 8 | -0.157 | 0.018 | 0.004 | 0.033 | 0.014 | 1.865 | 0.370 | 3.382 |
| Cv | soil_bulk_density_g_cm3 | 102 | 11 | -0.400 | 0.229 | 0.040 | 0.418 | 0.018 | 25.679 | 4.035 | 51.826 |

| | | | | | | | | | | | |
|---|---|---|---|---|---|---|---|---|---|---|---|
| Cv | usda_sand_percent | 113 | 12 | -0.174 | 0.002 | 0.000 | 0.003 | 0.028 | 0.162 | 0.018 | 0.307 |
| Cp | delta_bulk_density_percent | 54 | 5 | -0.086 | -0.010 | -0.013 | -0.007 | <0.001 | -0.997 | -1.265 | -0.728 |
| Cp | application_rate_t_ha | 27 | 4 | -0.069 | 0.003 | 0.002 | 0.003 | <0.001 | 0.259 | 0.172 | 0.346 |
| Cp | post_application_soil_bulk_density_g_cm3 | 54 | 5 | 0.402 | -0.289 | -0.454 | -0.125 | <0.001 | -25.109 | -36.468 | -11.719 |
| Cp | soil_porosity_percent | 27 | 4 | -1.234 | 0.024 | 0.010 | 0.038 | <0.001 | 2.428 | 1.015 | 3.860 |
| Cp | usda_sand_percent | 49 | 5 | 0.137 | -0.003 | -0.005 | -0.001 | 0.004 | -0.313 | -0.527 | -0.099 |
| Cp | usda_silt_percent | 58 | 6 | -0.147 | 0.004 | 0.001 | 0.007 | 0.012 | 0.372 | 0.081 | 0.663 |

Table S5. Estimated biochar effects by categorical moderator level. Estimates were obtained from three-level meta-analytic models without an intercept, allowing each retained category to have its own pooled effect estimate. Categories were retained only when represented by at least five effect sizes from at least two independent studies.

| thermal_property | moderator | level | estimate_lnrr | ci_lb_lnrr | ci_ub_lnrr | p_value | percent_change | ci_lb_percent | ci_ub_percent | n_effect_sizes_level | n_studies_level |
|---|---|---|---|---|---|---|---|---|---|---|---|
| k | feedstock_group | Agricultural Residues | -0.236 | -0.311 | -0.161 | <0.001 | -21.01 | -26.74 | -14.83 | 182 | 15 |
| k | feedstock_group | Non-food crop biomass | -0.106 | -0.224 | 0.011 | 0.076 | -10.10 | -20.07 | 1.10 | 9 | 2 |
| k | feedstock_group | Woody Biomass | -0.081 | -0.193 | 0.031 | 0.158 | -7.76 | -17.55 | 3.18 | 28 | 5 |
| k | climate_zone_by_lat_and_long | Subtropics | -0.266 | -0.364 | -0.167 | <0.001 | -23.32 | -30.51 | -15.39 | 89 | 5 |
| k | climate_zone_by_lat_and_long | Temperate | -0.139 | -0.205 | -0.073 | <0.001 | -12.99 | -18.54 | -7.07 | 100 | 13 |

| | | | | | | | | | | | |
|---|---|---|---|---|---|---|---|---|---|---|---|
| α | pyrolysis_temperature_group | <450 | -0.148 | -0.191 | -0.105 | <0.001 | -13.76 | -17.41 | -9.94 | 41 | 7 |
| α | pyrolysis_temperature_group | >600 | -0.058 | -0.125 | 0.008 | 0.086 | -5.65 | -11.72 | 0.83 | 7 | 2 |
| α | pyrolysis_temperature_group | 450–600 | -0.095 | -0.135 | -0.055 | <0.001 | -9.05 | -12.59 | -5.36 | 70 | 7 |
| Cv | soil_texture_group | Clay loam | -0.186 | -0.245 | -0.128 | <0.001 | -17.01 | -21.72 | -12.02 | 15 | 2 |
| Cv | soil_texture_group | Sandy loam | -0.101 | -0.149 | -0.052 | <0.001 | -9.58 | -13.86 | -5.09 | 34 | 4 |
| Cv | soil_texture_group | Silt loam | -0.002 | -0.050 | 0.046 | 0.933 | -0.21 | -4.89 | 4.71 | 48 | 4 |
| Cv | biochar_particle_size_group | Coarse | 0.076 | -0.026 | 0.178 | 0.142 | 7.91 | -2.52 | 19.46 | 10 | 2 |
| Cv | biochar_particle_size_group | Medium | -0.141 | -0.164 | -0.118 | <0.001 | -13.16 | -15.16 | -11.10 | 55 | 7 |
| Cv | feedstock_group | Agricultural Residues | -0.117 | -0.178 | -0.057 | <0.001 | -11.07 | -16.32 | -5.49 | 81 | 11 |
| Cv | feedstock_group | Non-food crop biomass | -0.033 | -0.110 | 0.044 | 0.403 | -3.24 | -10.42 | 4.52 | 9 | 2 |
| Cv | feedstock_group | Woody Biomass | 0.008 | -0.072 | 0.088 | 0.846 | 0.79 | -6.92 | 9.14 | 20 | 3 |
| Cv | climate_zone_by_lat_and_long | Subtropics | -0.151 | -0.212 | -0.089 | <0.001 | -14.01 | -19.14 | -8.55 | 44 | 4 |

| | | | | | | | | | | |
|---|---|---|---|---|---|---|---|---|---|---|
| Cv | climate_zone_by_lat_and_long | Temperate | -0.053 | -0.098 | -0.008 | 0.022 | -5.15 | -9.34 | -0.77 | 78 | 9 |

Table S6. Pairwise co-variation among biochar-induced changes in soil thermal properties. Pearson and Spearman correlations were calculated using log response ratios and pairwise complete observations. Pairwise sample sizes differ because not all studies reported all thermal properties under the same experimental contrasts.

| Property 1 | Property 2 | n | Pearson r | Pearson p | Spearman ρ | Spearman p |
|---|---|---|---|---|---|---|
| α | k | 86 | 0.819 | <0.001 | 0.743 | <0.001 |
| Cv | k | 81 | 0.799 | <0.001 | 0.784 | <0.001 |
| Cp | k | 52 | -0.013 | 0.926 | 0.264 | 0.058 |
| α | Cv | 80 | 0.571 | <0.001 | 0.578 | <0.001 |
| Cp | Cv | 46 | 0.431 | 0.003 | 0.533 | <0.001 |
| α | Cp | 52 | -0.146 | 0.300 | -0.084 | 0.555 |

### Supplementary Methods S4. Leave-one-study-out sensitivity analysis

Leave-one-study-out sensitivity analyses were conducted to assess whether pooled effects were driven by individual studies. For each thermal property, the three-level random-effects model was refitted after removing one study at a time. Stability was evaluated by comparing the direction, magnitude, confidence intervals, and p-values of the leave-one-study-out estimates with the main pooled estimate.

Table S7. Summary of leave-one-study-out sensitivity analyses for biochar effects on soil thermal properties. For each thermal property, the table reports the range of pooled percentage changes obtained after removing one study at a time, whether the direction of the pooled effect remained stable, and whether statistical significance was retained across all leave-one-study-out models.

| thermal_property | n_leave_one_out_models | min_percent_change | max_percent_change | main_direction_stable | all_significant_same_direction | min_p_value | max_p_value |
|---|---|---|---|---|---|---|---|
| Cp | 6 | -0.87 | 6.06 | FALSE | FALSE | 0.338 | 0.884 |

| | | | | | | | |
|---|---|---|---|---|---|---|---|
| Cv | 13 | -9.50 | -7.45 | TRUE | TRUE | <0.001 | 0.001 |
| α | 12 | -11.70 | -9.59 | TRUE | TRUE | <0.001 | <0.001 |
| k | 19 | -18.54 | -16.34 | TRUE | TRUE | <0.001 | <0.001 |

#### Supplementary Methods S5. Small-study effects and funnel asymmetry diagnostics

Small-study effects and funnel asymmetry were assessed for each thermal property using Egger-type meta-regression models fitted within the same three-level random-effects framework as the main analysis. The standard error of the effect size was included as a moderator of the log response ratio. A significant standard-error slope was interpreted as evidence of funnel asymmetry or small-study effects, but not as definitive evidence of publication bias because asymmetry can also arise from heterogeneity, study design, measurement conditions, and clustered effect sizes. Precision-based models using the inverse standard error were also fitted as a sensitivity diagnostic. Funnel plots were visually inspected for each thermal property (Figure S3).

Table S8. Small-study effects and funnel asymmetry diagnostics for biochar effects on soil thermal properties. Egger-type models used the standard error of the log response ratio as a moderator within the three-level random-effects framework. Precision-based models were fitted as sensitivity diagnostics. Significant slopes indicate potential small-study effects or funnel asymmetry, but should not be interpreted as definitive evidence of publication bias.

| thermal_property | n_effect_sizes | n_studies | main_lnr | main_percent_change | egger_slope_sei | egger_ci_lb | egger_ci_ub | egger_p | precision_slope | precision_p |
|---|---|---|---|---|---|---|---|---|---|---|
| k | 231 | 19 | -0.194 | -17.64 | -2.383 | -2.934 | -1.832 | <0.001 | 0.001 | <0.001 |
| α | 118 | 12 | -0.116 | -10.96 | -0.364 | -1.156 | 0.427 | 0.367 | 0.001 | 0.003 |
| Cv | 122 | 13 | -0.087 | -8.30 | 0.079 | -0.654 | 0.812 | 0.834 | 0.000 | 0.002 |
| Cp | 58 | 6 | 0.032 | 3.27 | 0.674 | -2.991 | 4.339 | 0.719 | 0.000 | 0.626 |

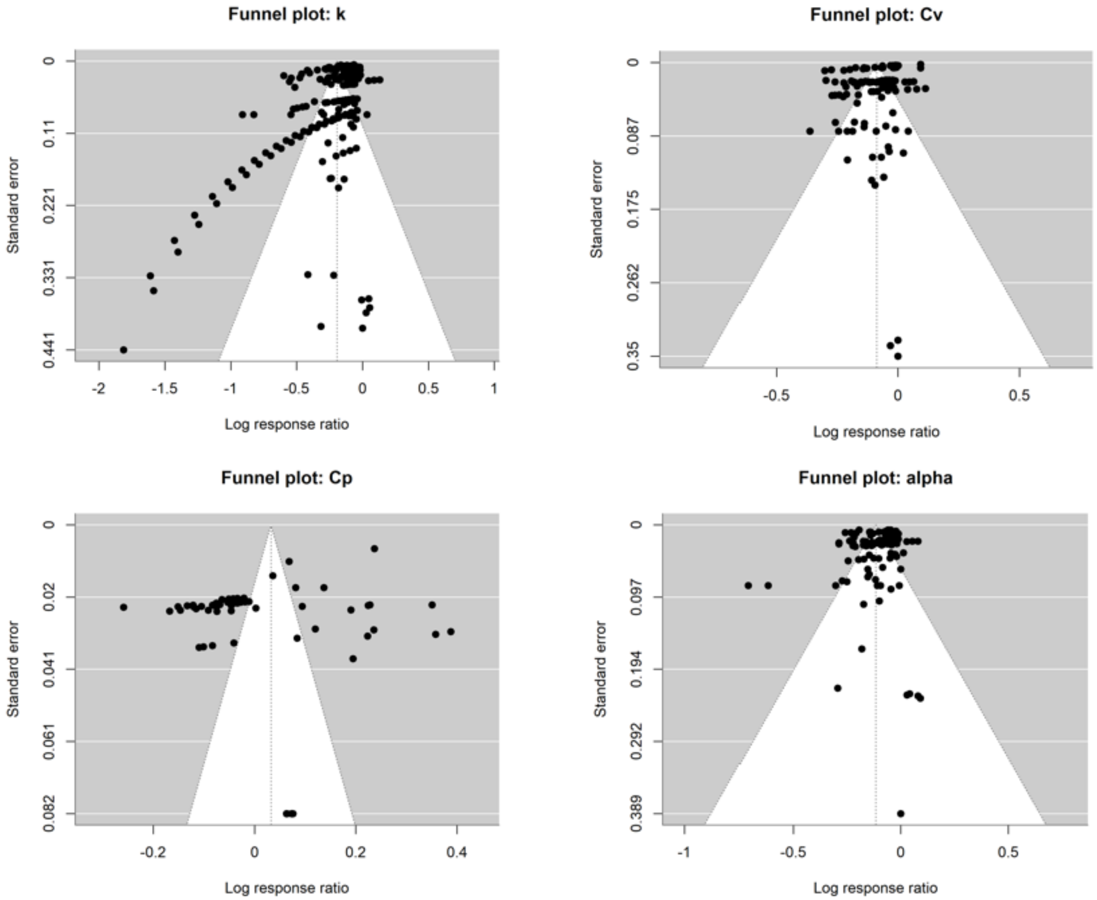


Figure S3. Funnel plots for biochar effects on soil thermal properties. Separate funnel plots are shown for thermal conductivity, thermal diffusivity, volumetric heat capacity, and gravimetric heat capacity. Plots are based on log response ratios and standard errors from the final cleaned effect-size dataset.
